# Transcriptomic analysis of quinoa reveals a group of germin-like proteins induced by *Trichoderma*

**DOI:** 10.1101/2021.04.14.439738

**Authors:** Oscar M. Rollano-Peñaloza, Patricia A. Mollinedo, Susanne Widell, Allan G. Rasmusson

## Abstract

Symbiotic strains of fungi in the genus *Trichoderma* affect growth and pathogen resistance of many plant species, but the interaction is not known in molecular detail. Here we describe the transcriptomic response of two cultivars of the crop *Chenopodium quinoa* to axenic co-cultivation with *Trichoderma harzianum* BOL-12 and *Trichoderma afroharzianum* T22. The response of *C. quinoa* roots to BOL-12 and T22 in the early phases of interaction was studied by RNA sequencing and RT-qPCR verification. Interaction with the two fungal strains induced partially overlapping gene expression responses. Comparing the two plant genotypes, a broad spectrum of putative quinoa defense genes were found activated in the cultivar Kurmi but not in the Real cultivar. In cultivar Kurmi, relatively small effects were observed for classical pathogen response pathways but instead a *C. quinoa*-specific clade of germin-like genes were activated. Germin-like genes were found to be more rapidly induced in cultivar Kurmi as compared to Real. The same germin-like genes were found to also be upregulated systemically in the leaves. No strong correlation was observed between any of the known hormone-mediated defense response pathways and any of the quinoa-*Trichoderma* interactions. The differences in responses are relevant for the capabilities of applying *Trichoderma* agents for crop protection of different cultivars of *C. quinoa*.

## Background

*Trichoderma* is a genus of ascomycete fungi widely studied for its versatile interactions with other organisms. *Trichoderma* can feed or parasitize on other fungi, bacteria, oomycetes and nematodes. Several species of *Trichoderma* are also symbionts with plants and can promote plant growth by several, yet so far only partially known mechanisms. Strains of several symbiotic species in the *Trichoderma harzianum* species complex (e.g. *Trichoderma afroharzianum* T22 (1), previously called *T. harzianum*) are used commercially because they can substantially improve yields of several species of crops. The strain T22 can enrich the soil nutrient availability to plants (2), and several species of *Trichoderma* have also been shown to enhance plant growth through volatile compound emission (3, 4) and stimulate plant systemic defense responses (5, 6). Nevertheless, plants do not always benefit from these interactions as described for some maize cultivar in field trials and lab experiments (7). Plant growth inhibition by *T. harzianum* has also been observed in axenic co-cultures with quinoa seedlings (8).

Quinoa (*Chenopodium quinoa* Willd.) is an emerging crop of great interest due to its nutritional values (9) and its resistance to hostile environmental conditions, especially salinity and drought (10, 11). Quinoa seeds are gluten-free, contain all essential amino acids and its composition (vitamins, antioxidants, fatty acids and minerals) is highly suitable for human nutrition (12). Quinoa has a high genetic diversity, e.g. more than 4,000 accessions have been registered by the Food and Agriculture Organization (13). The high genetic diversity of cultivars is the result of many years of selection by the indigenous people of the Andean Altiplano, where quinoa may have been domesticated 7,000 years ago by pre-Columbian cultures (14).

Quinoa agricultural yields can be boosted by *Trichoderma* application, as previously described (15). However, the outcome of plant-*Trichoderma* interactions is not always beneficial. Plant genotype-specific growth inhibition by commercially available *Trichoderma* strains have been reported for lentils (16), tomato (17) and maize (7). Thus, the incompatibility of particular plants with particular biocontrol strains can lead to undesired agricultural losses. Therefore, there is a need to understand the genotype-specific interaction mechanisms that determine whether plant growth is promoted or inhibited by biocontrol agents like T22.

In this work, we have studied the molecular response mechanisms of two *C. quinoa* cultivars that experienced plant growth inhibition when treated with *T. harzianum* BOL-12 and *T. afroharzianum* T22 in axenic co-cultures. The response of quinoa to BOL-12 and T22 in the early phases of interaction was studied by transcriptomic analysis and RT-qPCR verification. Overall, we observed that upon interaction with the two fungal strains, a broad spectrum of putative quinoa defense genes were activated in Kurmi but not in the Real cultivar.

## Methods

### Biological materials

Seeds of quinoa (*Chenopodium quinoa* Willd.) cultivars Maniqueña Real (*Real*) and Kurmi were kindly supplied by PROINPA (Quipaquipani, Bolivia). *Trichoderma afroharzianum*, Rifai, T22, anamorph ATCC 20847 (1) was purchased from the American Type Culture Collection (Manassas, VA, USA). *Trichoderma harzianum* BOL-12QD (BOL-12) was isolated and provided by the Instituto de Investigaciones Farmaco-bioquímicas (IIFB-UMSA, La Paz, Bolivia).

### Fungal growth

T22 and BOL-12 were maintained on potato dextrose agar (BD-Difco, Detroit, USA) at 25°C. To isolate conidiospore suspensions, one ml of sterile water was added to two-week-old *Trichoderma* cultures on potato dextrose agar and collected spores were filtered through a sterile piece of cotton wool. The spores were washed twice with sterile ddH_2_O and pelleted at 3700g for 5 min at 4 °C in an Allegra X-12R centrifuge (Beckman, Brea, CA, USA). Spores were resuspended in sterile ddH_2_O and kept at 4°C until experiments.

Germination of T22 and BOL-12 spores for *C. quinoa* treatment was performed as described by Yedidia, Benhamou (18) using 15 ml tubes shaken at 200 rpm for 18 h. The germinated spore suspension was washed twice by centrifugation as described above and finally resuspended in sterile ddH_2_O. The final spore concentration was adjusted to be 1 germinated spore/μl and verified by colony forming unit (CFU) counts in potato dextrose agar Petri dishes.

### Sterilization of *C. quinoa* seedlings and germination

Seeds of *C. quinoa* were surface-sterilized by soaking in commercial bleach (NaClO; 27 g/kg) for 20 min., followed by 6 rinses in sterile ddH_2_O. Immediately thereafter, the seeds were placed on sterile water agar (8 g/L) in Petri dishes and incubated in darkness at 24°C for 14 hours (8).

### Co-culture of quinoa and *T. harzianum* in Petri dishes

Five germinated axenic seedlings of each cultivar Kurmi and Real with similar root length were aligned on a straight line on 12×12 cm square Petri dishes containing 0.1X Murashige and Skoog Basal Salts Mixture (MS; Duchefa, Haarlem, The Netherlands), supplemented with 8 g/L agar. The Petri dishes were then tilted 45° during growth with the agar/air interface facing upwards and seedlings having the roots pointing towards the bottom part of the Petri dish. The seedlings were incubated at 24°C for 4 hours before treatment with T22 or BOL-12.

*C. quinoa* seedlings were treated by adding 10 μl [1 CFU/μl] of either T22 or BOL-12 germinated spore suspension on the neck of the primary root. Ten μl of sterile ddH_2_O was added to each seedling in the mock control group. After treatment, the seedlings were incubated at 24°C in a 16 h light /8 h dark photoperiod. Co-cultivation was done under fluorescent lights (Polylux XLr 30W, GE, Budapest, Hungary) at 50 μmol m^−2^ s^−1^ for 12 and 36 hours.

### Seedling growth analysis

Hypocotyl length was analyzed from images taken with a Digital Camera Canon EOS Rebel T3. Measurements from the photographs were done with the segmented line tool of *ImageJ* 1.49 (19).

### Sample collection and RNA extraction

For RNA extraction quinoa seedling were sampled 12 and 36 h after *Trichoderma* treatment as follows. The roots were excised at the root-hypocotyl interface with a scalpel. One root from each of five plates per treatment was pooled into one replicate on pre-weighed aluminum foil envelopes. The envelopes were weighed on a precision balance and shock-frozen in liquid nitrogen. Frozen samples were either processed immediately or stored at −80°C until RNA extraction. Roots and shoots were pooled separately.

Total RNA was extracted using the RNeasy Plant Mini Kit (Qiagen, Valencia, CA, USA), with the following modification: Root tissue samples preserved in liquid nitrogen were placed in a precooled mortar containing liquid nitrogen followed by thoroughly grinding without letting the samples thaw. Then 450 μL of Buffer RLT (Qiagen, Valencia, CA, USA) supplemented with B-mercaptoethanol (1%) was added. Grinding continued until samples thawed and were transferred to a 1.5 ml microcentrifuge tube. The rest of the procedure was followed according to Qiagen instructions. Total RNA quantity and quality was determined with a NanoDrop spectrophotometer. DNase treatment was performed with the DNA-*free* kit (Ambion, Carlsbad, CA, USA), following the instructions of the manufacturer. The integrity and quality of the RNA was determined as follows: 500 ng. of DNase-treated RNA were dissolved in 8 μl of sterile water and split it in two aliquots, one placed on ice and the other placed at 37°, both for 20 minutes, immediately 2 μl of loading buffer was added to each sample and together were loaded to an agarose gel (2%) stained with ethidium bromide. The gel was run at 80V for 30 min and visualized in an UV-transilluminator. Samples with sharp 18S and 28S rRNA and showing no evidence of degradation were retained.

### RNA-seq library construction and sequencing

Total RNA treated with DNase was sent to IGA technology services (IGA, Udine, Italy; http://www.igatechnology.com) for poly(A)^+^ mRNA purification, strand-specific cDNA synthesis, library construction (Truseq stranded mRNA-seq) and sequencing using a HiSeq2500 (Illumina Inc., San Diego, CA, USA) in paired-end mode with a read length of 125 bp. Raw sequences have been deposited at the National Center of Biotechnology Information NCBI under the Sequence Read Archive (SRA): SUB9370528.

### Transcriptomic analysis

RNA-seq reads were checked for quality by FastQC (v.0.9.0) and mapped on the quinoa genome “Kd” (Yasui et al., 2016) and to the QQ74 coastal genome (20) by Tophat2 (v.2.2.9). Transcript abundances were assessed with HTSeq (v.0.9.1) with “intersection-nonempty” mode. Genes that had a minimum of 1 read mapped in each of the samples considered for analysis were included. Gene expression levels were measured as counts per million (CPM) (21). Library size normalization was performed using the trimmed mean of *M*-values (TMM) within the R package edgeR (v.3.14.0) (22, 23). Differential gene expression analysis (treated vs mock-treated) was performed using edgeR with TMM normalized libraries (21) with a false discovery rate (FDR) of 5% (q < 0.05) (24).

### Functional annotation of differentially expressed genes

Gene ontology (GO) term enrichment for sets of differentially expressed genes were estimated with Argot2 through sequence function prediction (25). Singular enrichment analysis (SEA) for biological processes was performed with AgriGO v2.0 (26). The statistical test for SEA was Fisher’s exact test and for false discovery rate the Yekutieli method was applied (27).

### cDNA synthesis and gene expression by qRT-PCR analysis

Synthesis of cDNA was carried out with 500 ng of total RNA added to each 20 μl reaction of the RevertAid H Minus Reverse Transcription Kit (Thermo Scientific). The cDNA samples were stored at −20°C for downstream analysis. qRT-PCR of plant RNA was performed in a CFX384 Touch Real-Time PCR system (Bio-Rad, Hercules, CA, United States) using Maxima SYBR Green qPCR Master Mix (Thermo Scientific) supplemented with 0.25 μM of each specific primer and 10 ng of cDNA as template in a total volume of 10 μl/reaction. The PCR program had the following conditions: 1 cycle of: 95° C, 20 s; 30 cycles of: (95° C, 15 s; 60°C, 20s; 72 °C, 20s). The specificity of each PCR amplification was determined by melting curve analysis and by analysis in 2% agarose gels. The primer sequences can be found in Table S7. The relative transcript expression was calculated by the Pfaffl algorithm using, *CqACT2A* and *CqMON1*, as reference genes. Ten-fold dilutions of cDNA template were used to determine the amplification efficiency for each gene (28).

Primer pairs were designed using Perlprimer (29) so that one of the primers in each pair spanned an exon-exon border, and the primer pairs were additionally checked using Netprimer (https://premierbiosoft.com) to avoid primer-primer interactions.

### Protein evolution

Protein alignments were made using Muscle (30). Aminoacid substitution models were evaluated by MEGA X (31). The protein evolutionary tree was performed by maximum likelihood using LG model (32) with gamma distribution (LG + G) and 95% limit for partial gaps. Total positions in the final dataset were 191. Bootstrap testing was conducted with 1000 replicates.

## Results

### *Trichoderma* BOL-12 and T22 inhibit the growth of *C. quinoa* seedlings under certain axenic conditions

Previously we have observed that quinoa growth was inhibited by two *Trichoderma* strains under axenic conditions (8). Therefore, we decided to investigate the axenic co-culture of quinoa with T22 and BOL-12 by gene expression analysis to detect the molecular signaling possibly responsible of the *C. quinoa* growth inhibition. Briefly, quinoa seedlings of cultivars Kurmi and Real were grown for 18 h in square petri dishes on 0.1x MS and 0.8% agar and co-cultivated with T22 or BOL-12 for 12 and 36 hours. The studied *Trichoderma* strains did not have any measurable effect on the growth of *C. quinoa* seedlings during this short time of co-cultivation. Thus, the growth pattern was consistent with previously reported.

### Transcriptome sequencing of *C. quinoa* in axenic co-culture with *Trichoderma*

RNA samples from quinoa roots treated with *Trichoderma* for 12 and 36 hours were collected. RNA samples at 12 hours post inoculation (hpi) were analysed through RNA-seq and RNA samples at 36 hpi were evaluated through posterior qRT-PCR. For the transcriptomic analysis three biological replicates of each treatment were sequenced in paired-end mode. The final number of reads that passed the quality control varied between 10.2 and 23.1 million paired-end reads of 125 bp per sample (Table 1). Reads were mapped to the draft quinoa genome of cultivar Kd as well as to the chromosome-level assembly of the quinoa genome cultivar QQ74 (Table 1). On average, the proportion of mapped reads was substantially increased when reads were mapped to the QQ74 quinoa genome (93.9 %), as compared to the draft Kd quinoa genome (71.8%). Therefore, all downstream analyses were performed with data mapped to the QQ74 genome.

**Table 1.**
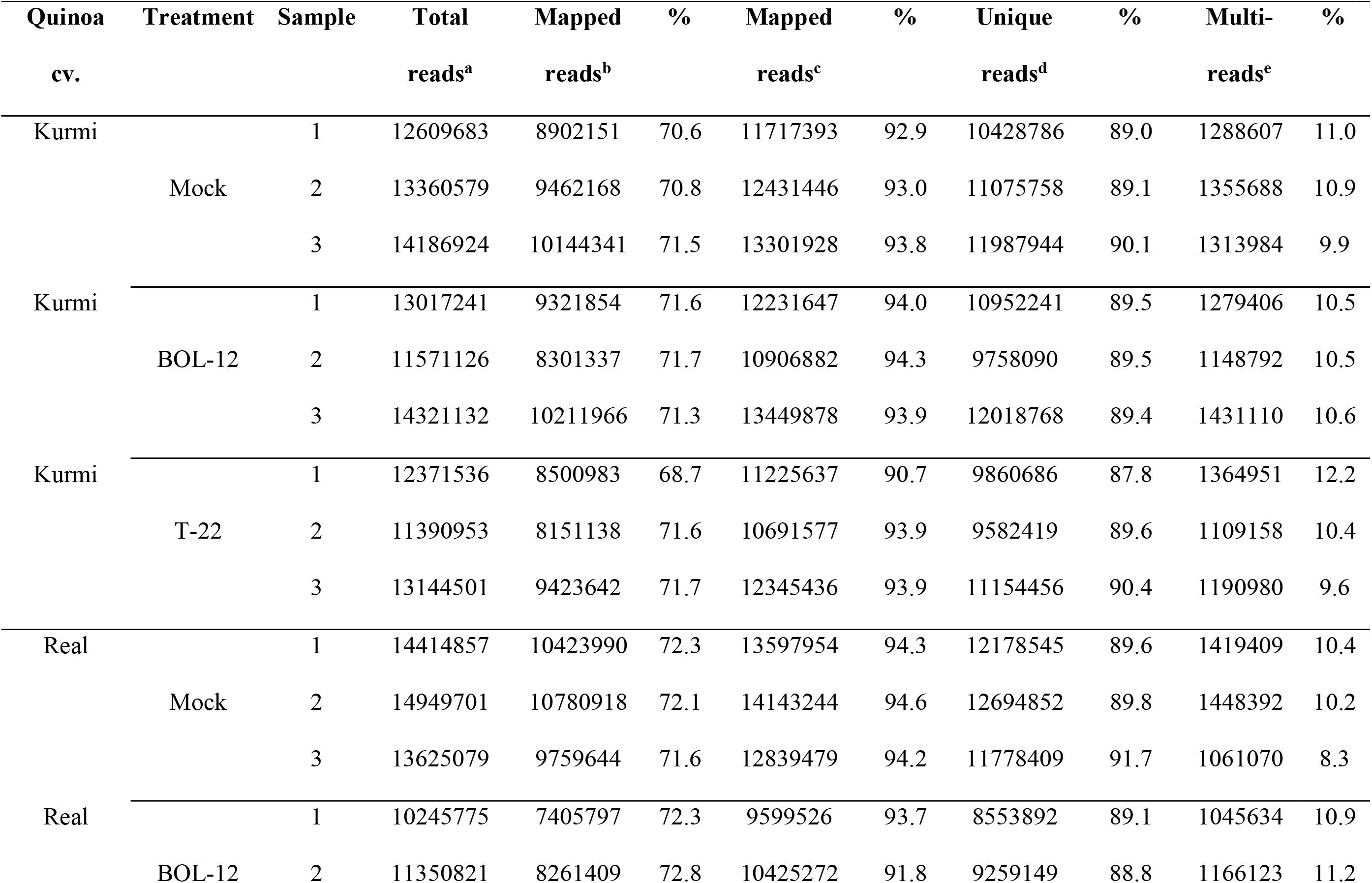
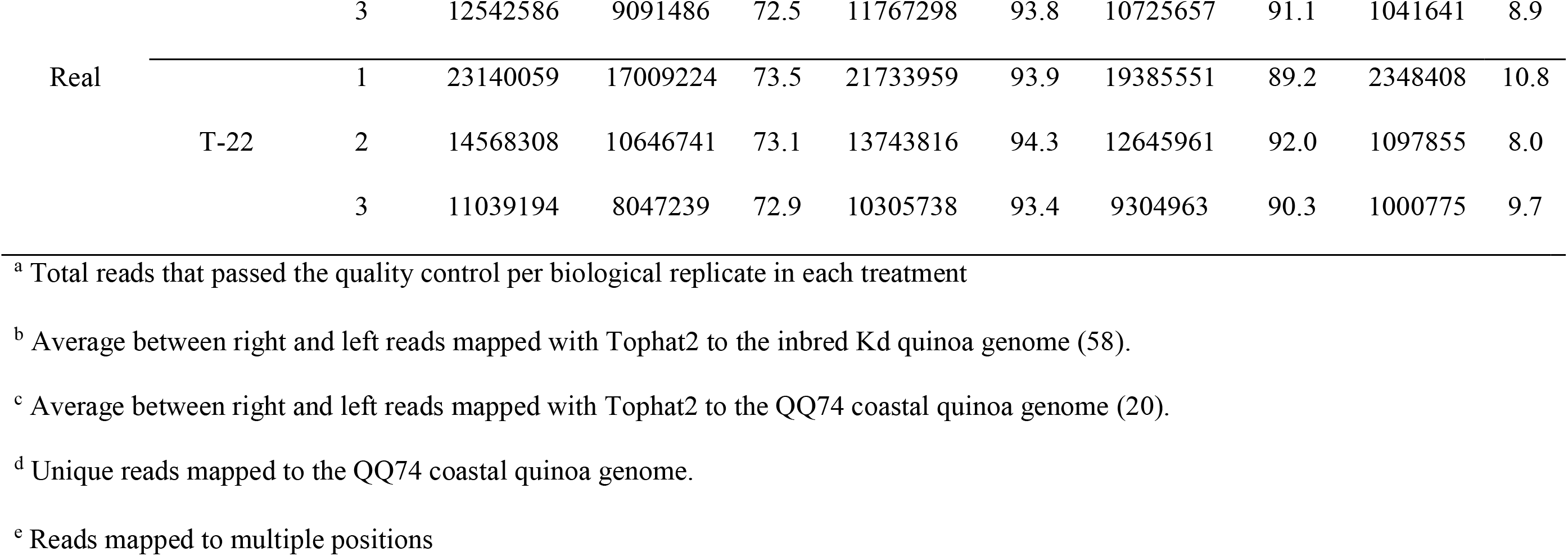
RNA-seq summary of read numbers mapped to two quinoa reference genomes

### Differential gene expression in quinoa in response to *Trichoderma* treatment

The differential gene expression analysis considered only reads that mapped to unique locations in the QQ74 genome. The average number of reads that were mapped to unique locations in the QQ74 genome was 89.8% (Table 1). The remaining reads (10.2%) producing multiple alignments were discarded. Further, only quinoa genes with at least one read in each of the samples analysed were considered (Table 2). Gene expression levels were measured as counts per million (CPM). CPM were TMM-normalized in order to compensate for library size differences. Differential gene expression analysis comparing mock-treated samples with samples treated with *Trichoderma* was performed with edgeR.

**Table 2.** Differentially expressed genes in quinoa roots treated with *Trichoderma*

| Quinoa | Experiment | Induced | Repressed | Total | Ind/Repr | Genes evaluated<sup>a</sup> |
| --- | --- | --- | --- | --- | --- | --- |
| Kurmi | BOL-12 vs. mock-treated | 158 | 38 | 196 | 4.2 | 25 273 |
| Kurmi | T22 vs. mock-treated | 1417 | 1727 | 3144 | **0.8** | 25 379 |
| Real | BOL-12 vs. mock-treated | 277 | 76 | 353 | 3.6 | 30 108 |
| Real | T22 vs. mock-treated | 1170 | 139 | 1309 | **8.4** | 30 745 |

<sup>a</sup> Genes included had at least one read in each of the samples. The quinoa genome annotation contains 44 776 genes.

Quinoa roots in general induced more genes than they repressed upon interaction with *Trichoderma*, with the exception of Kurmi interacting with T22 where more genes were repressed than induced (Table 2). Kurmi treated with T22 showed 16 times more differentially expressed genes than in the treatment with BOL-12. Similarly, quinoa cv. Real treated with T22, compared to the mock-controls, had 5.5 times more differentially expressed genes than Kurmi in the treatment with BOL-12 (Table 2).

Regarding communal effects by both *Trichoderma* strains, we observed more genes differentially expressed in cv. Real (141 genes) than in cv. Kurmi (75 genes) (Tables S1-3). Among the quinoa genes up- or downregulated under one or several conditions, only 19 were communally differentially expressed in all experimental combinations, and all were induced (Figure 1, Table 3). That is, they were significantly induced during the interaction of each quinoa cultivar with each *Trichoderma* strain. The group of 19 differentially expressed genes were not significantly associated with any functional GO term upon analysis by SEA. However, the GO analysis suggests that 13 of the 19 gene products are localized outside the cytoplasm. This indicates activity at the plasma membrane, the cell wall and the extracellular compartment, indicating functions relating to interactions with external stimuli (Table 3).

**Table 3.**
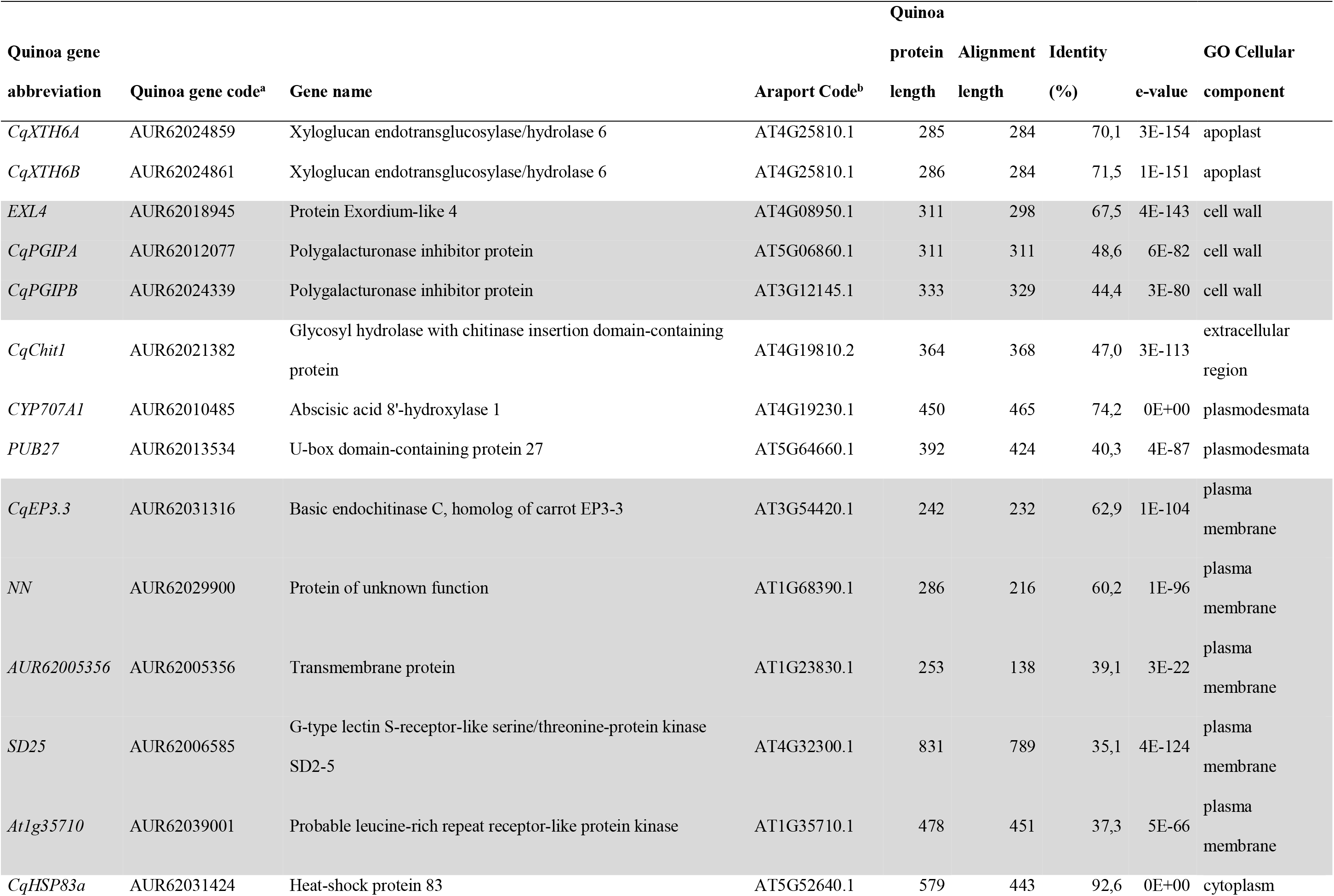
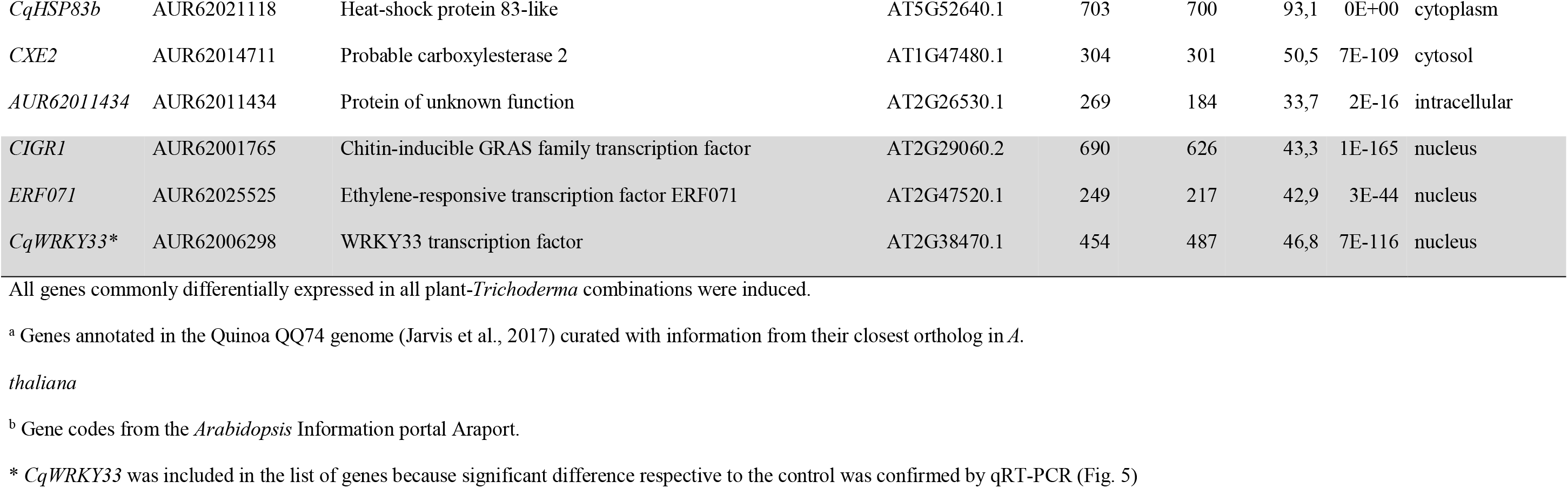
Quinoa genes differentially expressed in both Kurmi and Real in response to both *Trichoderma* strains

**Figure 1.**
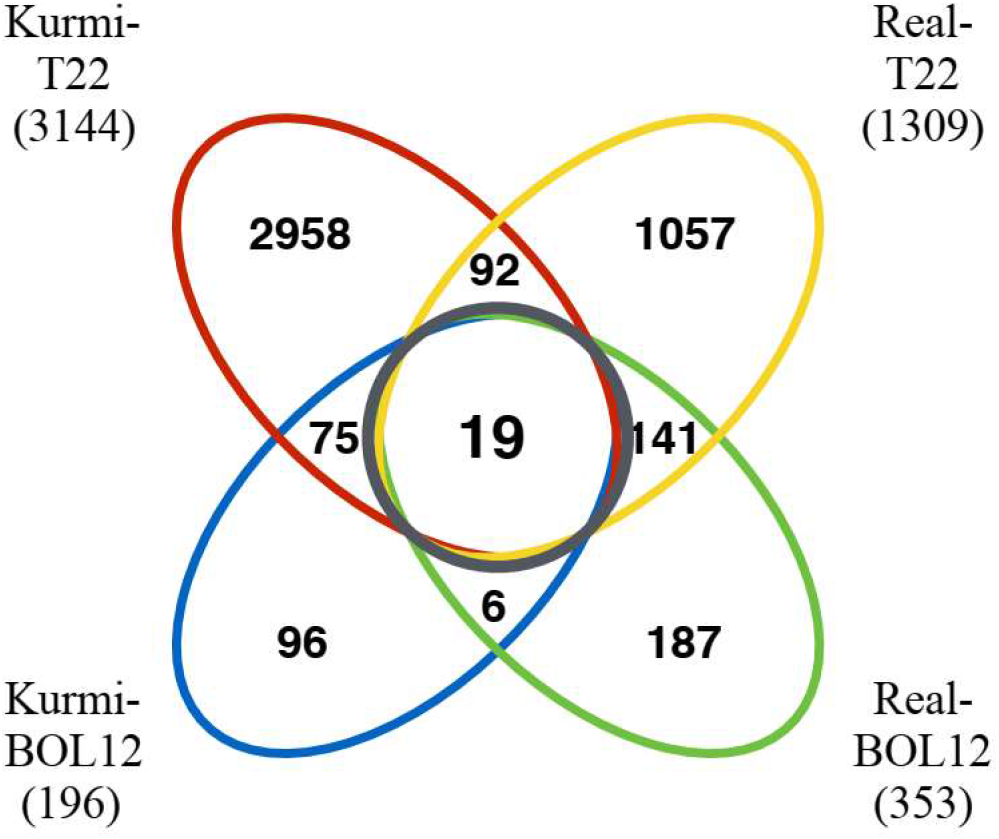
Venn diagram of quinoa genes differentially expressed in response to *Trichoderma*. Quinoa genes differentially expressed were grouped according to the cultivars and *Trichoderma* strains studied. The black circle indicates genes differentially expressed in both quinoa cultivars by each of the *Trichoderma* strains tested. The numbers in parenthesis indicate the number of genes differentially expressed in each of the quinoa-*Trichoderma* interactions studied.

### Quinoa genes differentially expressed unique to each cultivar

We decided to analyze genes that were induced by *Trichoderma* and were uniquely expressed in each cultivar. The Kurmi cultivar upon interaction with either BOL-12 or T22, expressed 75 genes that were communally differentially expressed (DE) but were not differentially expressed upon either *Trichoderma* interaction in cv. Real (Figure 1 and 2, Table S1). The expression profiles of these genes were clustered by Euclidean distance and are shown in Figure 2. From the 75 DE genes in cv. Kurmi by both strains of *Trichoderma*, 59 genes were induced (Table S1), whereas 16 DE genes were repressed (Table S2). The 75 DE genes expressed in cv. Kurmi are expressed in cv. Real but are not responsive to the treatment with either of the *Trichoderma* strains (Figure 2).

**Figure 2.**
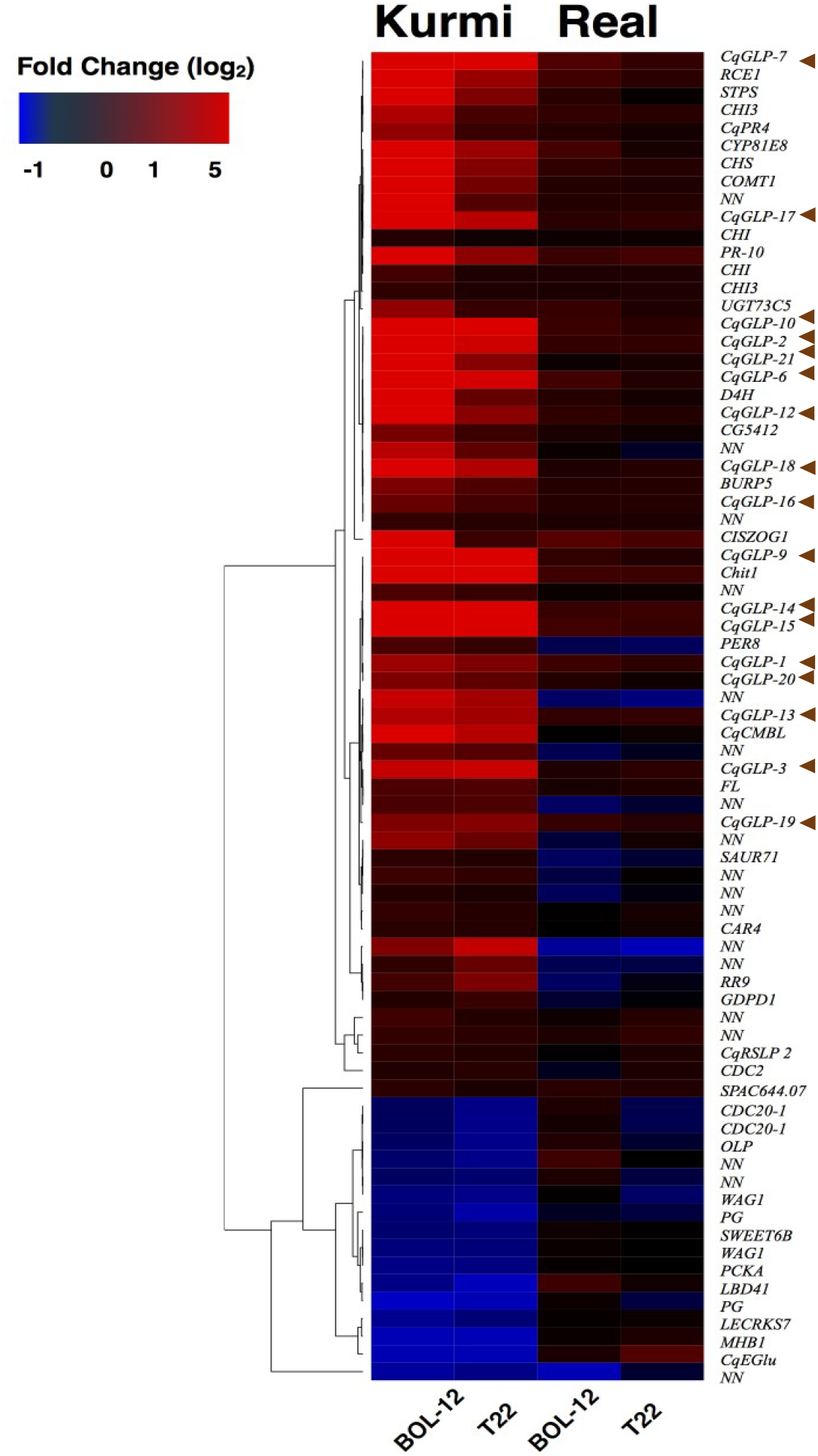
Heatmap profile of genes responsive to BOL-12 and T22 in Kurmi but not in Real. Genes differentially expressed in cv. Kurmi in response to either *Trichoderma* strain, but not significantly responsive in cv. Real, were analysed. Clustering by Euclidean distance shows the similarity in expressional change upon *Trichoderma* treatment. Brown arrows indicate *CqGLPs*.

Analysis of the 59 significantly induced genes revealed that 17 genes (*CqGLPs*) are highly expressed and share a high protein sequence identity (90%). These genes encode proteins that belong to the germin-like protein family (GLPs) (Figure 2; Table S1). Further, several genes involved in flavonoid biosynthesis were specifically responsive in Kurmi. We identified 9 genes whose orthologs in *Arabidopsis thaliana* are described to be involved in the flavonoid biosynthesis pathway. These differentially expressed genes are orthologs to four out of five enzyme-coding genes necessary for production of flavonol glycosides from naringenin, also known as chalcone (33) (Figure 2, Table S1).

The Real cultivar had 141 genes differentially expressed common to both *Trichoderma* strains tested (Figure 1, Table S3). The cv. Real response to both *Trichoderma* strains showed mostly activation of transcription factors and enzymes without a significant match to a known pathway. Among the genes that were differentially expressed there are 4 ethylene-responsive transcription factors, 9 probable WRKY transcription factors and 3 chitinases (Table S3).

Genes differentially expressed related to biotic interactions were observed in mayor proportions in quinoa cv. Kurmi than Real. Therefore, the focus of this study was on the response of the Kurmi cultivar.

### Functional annotation of differentially expressed genes

To assess the function of the differentially expressed genes we annotated the differentially expressed genes of all combinations with Gene Ontology (GO) terms for biological processes. The quinoa genome has 44 776 annotated genes (20) but the annotation with Argot only assigned GO terms to 50.5 % of the genes (i.e. 22 650 genes annotated with GO terms). Despite the low percentage of GO terms assigned, GO annotation for the biological process category in Kurmi plants treated with BOL-12 revealed defense response (GO:0006952) and response to biotic stimulus (GO:0009607) as the main and only processes associated to *Trichoderma* BOL-12 treatment (Table S4). In contrast, the interaction between Kurmi and T22 did not show any significant GO term for biological processes.

Quinoa plants of the Real cultivar had more genes associated to GO terms than Kurmi. However, no specific association to a cluster of similar GO terms were observed (Table S4). Specifically, Real treated with T22 showed 38 genes that were annotated to response to stress (GO:0006950) and 6 genes that were annotated to chitin catabolic processes (GO:0006032) and associated redundant GO terms (Table S4). Further, Real treated with BOL- 12 did not show any GO terms directly associated to defense response or response to stress, yet highest significance was observed to GO terms for cell wall related processes (Table S4). Nonetheless, in the interaction between Real and each strain of *Trichoderma* we observed several genes related to defense being commonly activated. Among them WRKY transcription factors (9 differentially expressed genes), ethylene-responsive genes (4 differentially expressed genes) and chitinases (3 differentially expressed genes) (Table S3).

### Validation of RNA-seq with qRT-PCR

Quinoa root transcriptomes have not previously been analysed. We therefore validated the gene expression data obtained by RNA-seq by performing qRT-PCR for 10 selected genes, including induced, repressed and stably expressed genes (Figure 3, Table S5). A log-log linear model analysis of the RNA-seq data and the qRT-PCR data showed a strong correlation (R^2^) of 0.848. The correlation was higher when the different quinoa-*Trichoderma* interaction pairs were assessed independently (Table S5).

**Figure 3.**
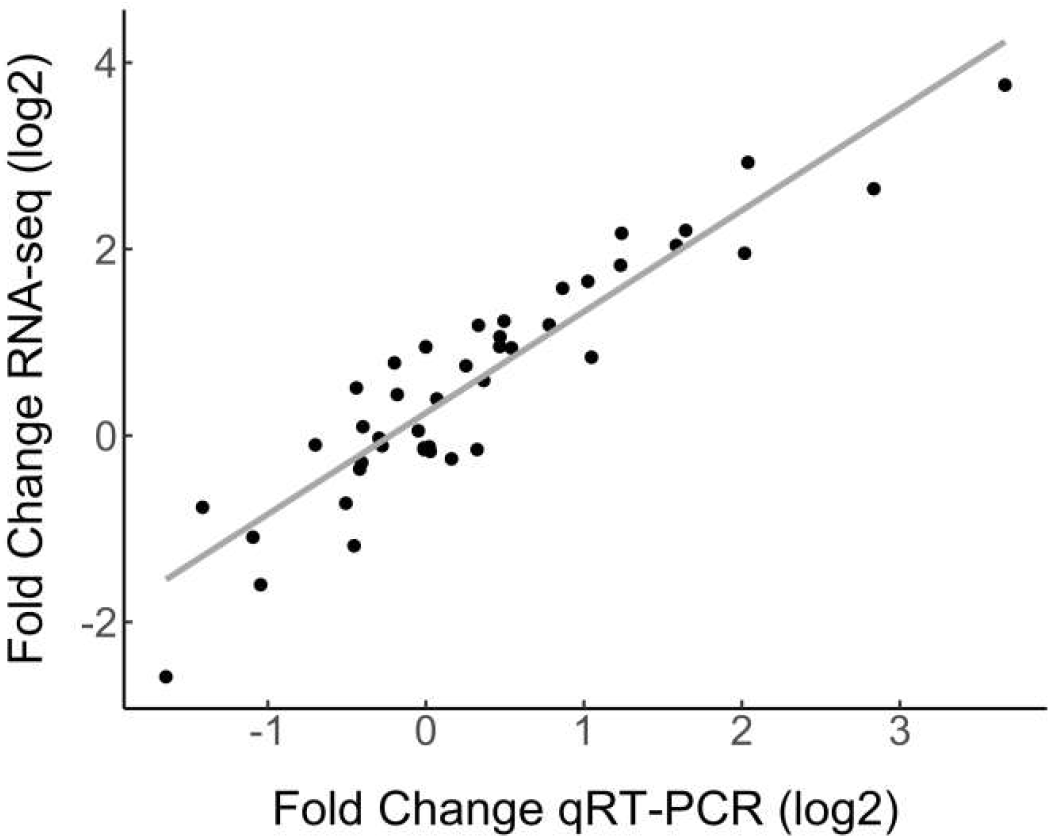
Correlation of RNA-seq and qRT-PCR gene expression data. Ten differentially expressed genes and two reference genes from the RNA-seq dataset were evaluated by qRT-PCR. Gene expression by qRT-PCR was normalized to the *CqAct2* reference gene. Fold change was measured by comparing samples treated with *Trichoderma* against mock- treated. The selected genes were assessed in all quinoa-*Trichoderma* combinations as averages of triple biological replicates. The Pearson correlation coefficient between the RNA-seq and qRT-PCR data was 0.921 (For data see Table S5).

### Changes in root gene expression at 36 hpi

Time changes in the expression of quinoa genes by *Trichoderma* treatment were assessed by qRT-PCR at 36 hpi. We followed time-dependent changes in two highly induced genes (*CqGLP1* and *CqGLP10*) representing the GLP family and one gene that was induced in all quinoa-*Trichoderma* interactions (*CqHSP83).* The gene expression of *CqGLP1* and *CqGLP10* was reduced at 36 hpi compared to 12 hpi in the Kurmi cultivar but its expression was still higher than the mock-treatment. In contrast, the gene expression of *CqGLP1* and *CqGLP10* in the Real cultivar was higher at 36 hpi than at 12 hpi, being statistically significant in the Real - BOL-12 interaction (Figure 4). The *CqHSP83* gene maintained its level of gene expression between 12 hpi and 36 hpi by application of T22 in both cultivars. In contrast, the application of BOL-12 to both cultivars downregulated *CqHSP83* gene expression at 36 hpi as compared to 12 hpi. However, the downregulation was only significant in the Kurmi cultivar (Figure 4). Overall, the results indicate that the induction of the analysed genes is slower in cv. Real than in cv. Kurmi.

**Figure 4.**
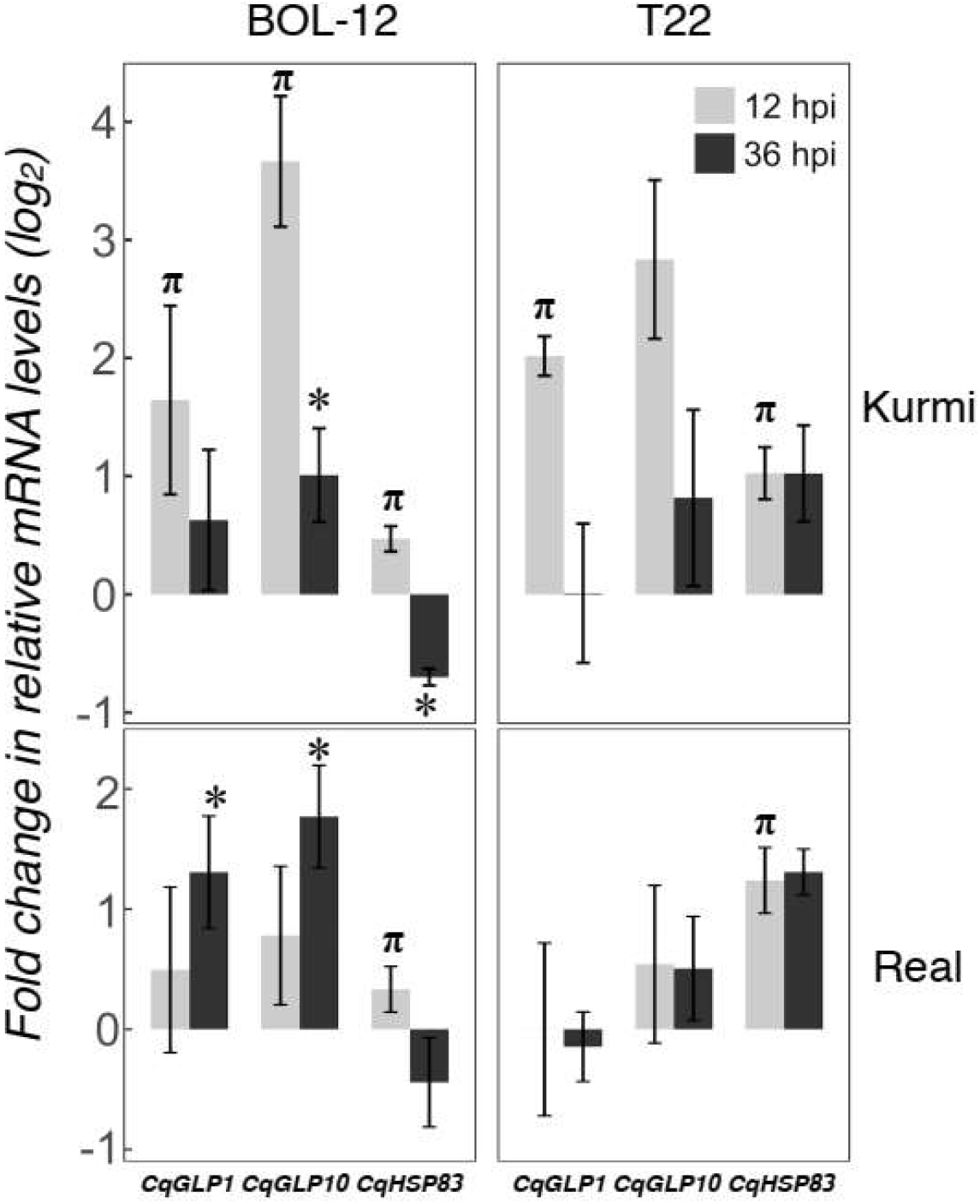
Gene expression changes at 12 and 36 hpi in representative *CqGLPs* and *CqHSP83*. Quinoa root samples treated with *Trichoderma* were assessed at 36 hpi by qRT-PCR. mRNA levels were normalized to the *CqAct2* reference gene. Fold changes (mean ± SE) were determined by comparing samples treated with *Trichoderma* against mock-treated ones. Asterisks denote significant changes in the gene expression as compared to the control treatment by qRT-PCR at 36 hpi (*p* < 0.05). Symbol π denotes significant changes in the gene expression as compared to the control treatment, using RNA-seq (*p* < 0.05) and confirmed by qRT-PCR (*p* < 0.05) at 12 hpi.

### Shoot gene expression

We investigated changes in the quinoa shoot gene expression 36 h after *Trichoderma* treatment at the root neck (Figure 5). Ten out of 12 genes investigated were also expressed in the shoots, *CqPER39* and *CqPR1C* gene expression was not detected at the shoots in any of the combinations studied (Table S6). *Trichoderma*-induced gene expression changes at the shoots (Figure 5) showed a generally similar pattern of gene expression as observed in the roots at 36 hpi (Figure 4). *CqGLP1* and *CqGLP10* are significantly expressed in both cultivars upon interaction with BOL-12 but not with T22. Likewise, *CqHSP83* is significantly expressed in both cultivars when interacting with T22 but not when interacting with BOL-12 (Figure 5). The other genes did not show a significant correlation in the Shoot-root expression in any of the quinoa-*Trichoderma* interactions studied (Table S6).

**Figure 5.**
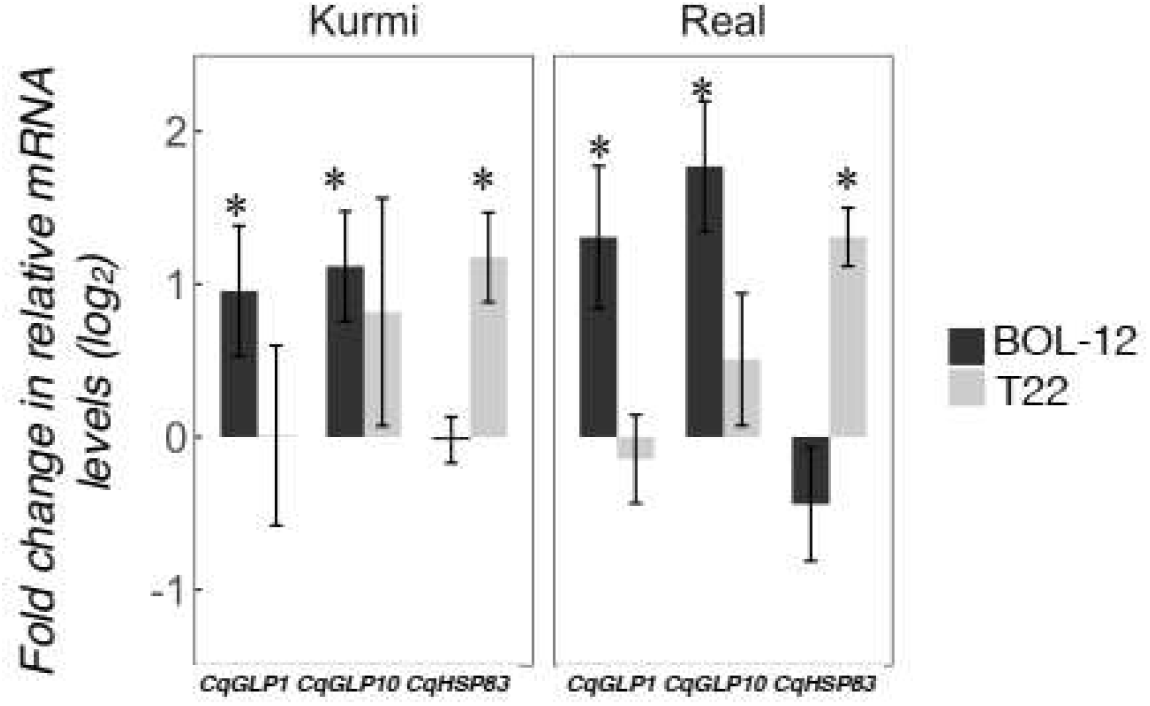
Gene expression changes in quinoa shoots after treatment of roots with *Trichoderma*. Quinoa shoot samples were assessed by qRT-PCR 36 h after treatment with *Trichoderma* to the roots. Gene expression was normalized to the *CqAct2* reference gene and is shown as mean ± SE. Asterisks denote significant changes in the gene expression as compared to the control treatment (*p* < 0.05). Two *CqGLP* genes that were induced in Kurmi roots at 12 hpi as well as the *CqHSP83* gene, which was induced in roots in all quinoa-*Trichoderma* interactions at 12 hpi are shown.

### Evolutionary analysis of the germin-like proteins

Plant germins were first investigated and have been characterised in most functional detail in cereals (34). To investigate the coincidental induction of germin-like proteins in cv. Kurmi (Table S1), we carried out BLAST searches to identify all germin and germin-like homologues in *C. quinoa*, *Beta vulgaris*, *A. thaliana* and *Hordeum vulgare* and performed alignments and evolutionary analyses. We found that 16 of the17 quinoa GLPs induced by *Trichoderma* (highlighted in green) in cv. Kurmi belong to a single (98% bootstrap) *C. quinoa*-specific clade of 29 homologues (Figure 6 and Table S8). The remaining GLP induced by *Trichoderma* (*CqGLP20*) groups in an unresolved putative clade, which contains four quinoa GLPs and one sugarbeet GLP (Figure 6). The relation of the quinoa-specific clade to homologues in other species, including the closest relative *B. vulgaris*, was not resolved, whereas other groups of quinoa germin-like proteins were significantly associated with specific homologues in *B. vulgaris.* Species-specific gene groups were also observed for *B. vulgaris* and *A. thaliana*. The result suggests that recent expansions of gene groups have occurred independently in the amaranth family species *C. quinoa* and *B. vulgaris*.

**Figure 6.**
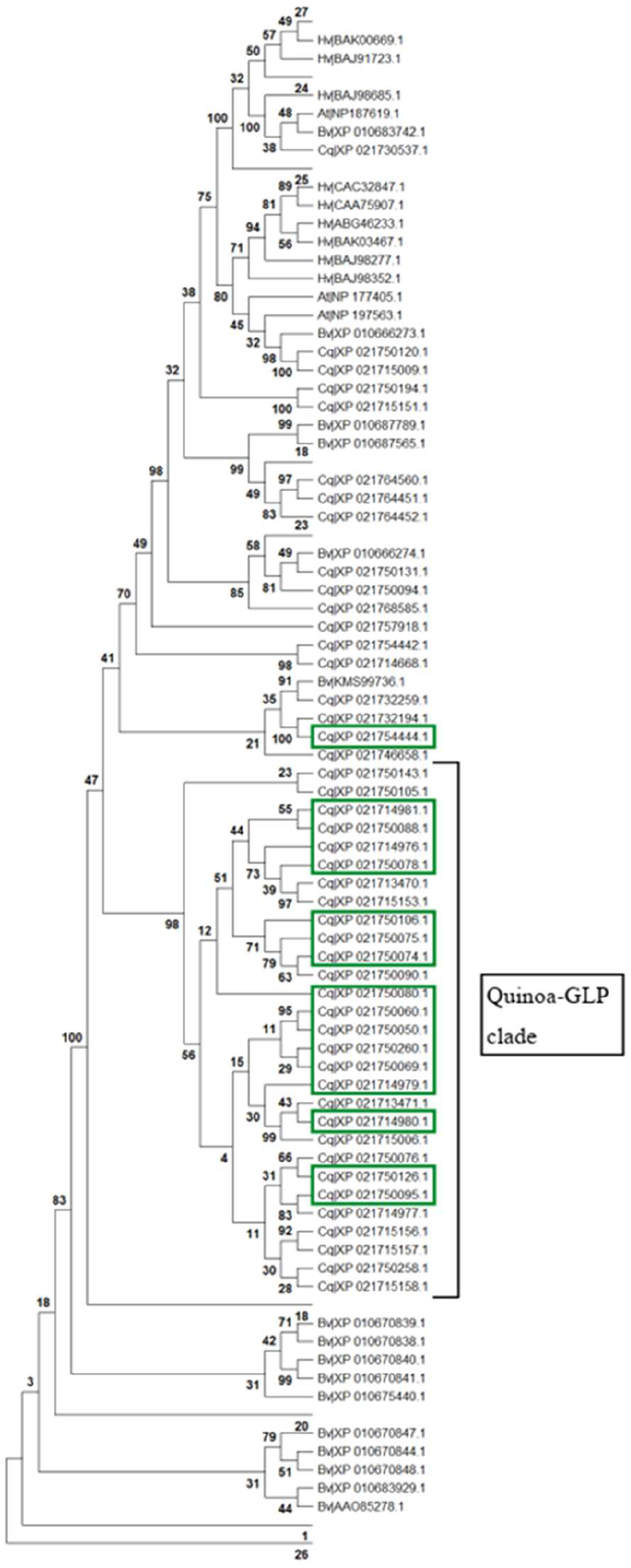
Protein evolutionary tree of germin and germin-like proteins. All identified germin-like proteins found in *C. quinoa* (Cq), *B. vulgaris* (Bv), *A. thaliana* (At) and *H. vulgare* (Hv) homologues were aligned by Muscle. The protein evolutionary tree was constructed by maximum likelihood using the LG + G model with 1000 iterations. Bootstrap values are given in percentage (%). Values below 30% are not shown except for in the Trichoderma-responsive quinoa-specific GLP clade. The *Trichoderma*-induced homologues are marked in green.

## Discussion

The outcome of plant-*Trichoderma* interactions with respect to both growth and physiological changes has been shown to be genotype-specific regarding both the plant and the *Trichoderma* biomaterial (5, 17). Here, we have observed a small set of quinoa genes being responsive in all combinations of *Trichoderma* strains and quinoa cultivars studied. However, we have found many more genes that are differentially expressed by a specific quinoa cultivar in response to either or both of two *Trichoderma* strains (Figure 1). Nonetheless, the outcome of the interaction of *Trichoderma* on plant growth in axenic co-cultures is negative for plants, similarly for both quinoa cultivars and consistent with previous observations (8).

The set of 19 genes that showed significant responses in all cultivar-strain combinations mainly include genes connected to biotic stress response and cell wall modification (Table 3). Orthologs of these genes are known to be involved in defense response. For example, the polygalacturonase inhibitor protein *AtPGIP1* (AT5G06860) in *A. thaliana* is thought to inhibit cell wall pectin degrading enzymes, commonly produced by fungal pathogens (35). Xyloglucan endotransglucosylase/hydrolases are cell wall repairing enzymes, many of which are induced by fungal infection (36). Further, two highly similar (protein sequence identity of 94,3%) heat-shock proteins (*CqHSP83A* and *CqHSP83B*) annotated to be involved in general stress responses (GO:0006950) were upregulated in all quinoa-*Trichoderma* interactions. Large heat-shock proteins (70 – 90 kDa) are known to be involved in plant defense response through the stabilization of protective plant proteins (37–39). The heat-shock proteins expressed by quinoa might have been induced to contribute to the stabilization of defense proteins that would prevent or counteract damages induced by *Trichoderma* (8).

Several differences in the defense response patterns between the quinoa cultivars were observed. Especially, the Kurmi cultivar that displays specific activation of several homologs to biotic stress-associated plant genes (Table S1-2). In contrast, the responsive genes in cv. Real were mostly involved in general cellular processes (Table S3), and to a lesser extent involving defense response genes. The defense response gene set induced in cv. Real was also completely different from the one activated in Kurmi (Figure 1, Table 2 and Table S1-3). Surprisingly, in neither case an obvious association to known major pathogen response pathways like jasmonic acid, salicylic acid or ethylene pathways (40–42) could be observed in the GO analysis at 12 hpi. The low association of the quinoa differentially expressed genes with these known pathways could, however, be caused by the relatively low level of GO annotation observed for the quinoa genome.

The quinoa genes specifically induced in the Kurmi cultivar upon interaction with *Trichoderma* (Table S1) resemble a set of defense response genes observed in plant-pathogenic interactions (43). Several of these induced genes (chalcone synthase, chalcone isomerase, flavonol synthase, UDP glycosyl transferase and *cis*-zeatin O-glucosyltransferase) belong to the flavonoid biosynthetic pathway (33). Flavonoids have an important role in plant defense (43, 44). For example, some *A. thaliana* mutants lacking the UDP glycosyl transferase gene (*AtUGT74F1*) are more susceptible to *Pseudomonas syringae* infection than the wild-type (45). Further, a QTL analysis for pathogen resistance in soybean identified two UDP glycosyl transferase genes as the candidate genes responsible for resistance to *Fusarium* (46). Thus, the Kurmi cultivar might be producing flavonol glycosides in order to prevent damage from *Trichoderma* overgrowth.

The Kurmi cultivar specifically induced several plant defensins that belongs to the germin and GLP family (Figure 6 and Table S1). The *Trichoderma*-responsive GLPs form a majority (16 genes out of 29) of a recently expanded quinoa-specific clade (Figure 6, Table S8), which are thus strongly connected to *Trichoderma* interaction. Two of the GLP genes were further tested, and were found to be also induced in leaves (Figure 5). The timing of the induction further indicated that GLPs are induced in both Kurmi and Real, albeit more slowly in Real (Figure 4). Especially studied in grains like barley, GLPs are plant proteins involved in defense response and characterized by various enzymatic activities including oxalate oxidase (OXO), superoxide dismutase (SOD), ADP-glucose pyrophosphatase/ phosphodiesterase (AGPPase) and polyphenol oxidase (PPO) (47, 48). A potential function of germin-like proteins (GLPs) is found in its OXO and SOD activities, which may play a key role in production of hydrogen peroxide (H_2_O_2_) during plant defense (34). Because of a potential major importance of GLP in protecting cells from superoxide toxicity produced under pathogen attacks, germin or GLP genes (i.e. *HvOXO1*) have been inserted into dicot plants like rapeseed or peanut to enhance its pathogen resistance (48–50). Given that several quinoa GLPs were expressed upon interaction with *Trichoderma*, it is very likely that these GLP defensins have an important role in the plant immune response of quinoa. Resistance to microbe attacks has been previously connected to the speed of response in different cultivars (51, 52). Thus, the rapid induction of a cluster of GLPs in both roots and leaves of Kurmi compared to Real (Table S2-3, Figure 4) makes these genes potential candidates for breeding to increase the tolerance to microbial attacks in quinoa plants.

The set of defense-related genes (peroxidases, chitinases, ERF and WRKY transcription factors) induced in quinoa cv. Real upon interaction with *Trichoderma* (Table S3) has been observed in *L. japonicus* roots upon 1 hour of incubation with chitin oligosaccharides (53). However, the levels of such defense-related genes returned to normal after 7 hours in *L. japonicus* whereas in quinoa remained induced after 12 hours, possibly due to the persistance of interaction with the living *Trichoderma* agent as compared to the transient nature of the elicitor. Similar to the *L. japonicus system*, the 24 h interaction of *Trichoderma* with *A. thaliana* resulted in the induction of the same defense-related genes as seen here in quinoa (54). This set of genes could thus be a basal gene response of plants after recognition of beneficial fungi like *Trichoderma* through chitin. In contrast, the Kurmi cultivar might have a different set of receptors that helps the plant to perceive a possible negative effect from *Trichoderma* and thus rapidly activate a different set of defense-related genes.

The plant root response to *Trichoderma* at transcriptomic level has been poorly studied compared to aerial parts (55). Nevertheless, it has been observed that in tomato roots the recognition of *Trichoderma* at 24 hpi activates ROS signaling, SA responses, cell wall modifications (56), JA responses and induction of plant defenses (54, 57). In our study, we have observed a similar pattern for ROS signaling, cell wall modifications and induction of plant defenses (Table 3, Table S1-3), confirming that the first response of root plants to beneficial fungi like *Trichoderma* is to activate defenses. Further, our study reveals that the defense response against beneficial fungi is variable between cultivars (Figure 2). The variable molecular response between cultivars to *Trichoderma*, could help to create molecular markers of compatibility between certain plant cultivars and certain strains of *Trichoderma*.

In conclusion, our study suggests that *Trichoderma* triggers a defense response in quinoa plants. Comparing the defense response of two quinoa cultivars we can observe that the Kurmi cultivar mainly induced a set of genes involved in plant defense. In contrast, the Real cultivar did not have a clear response because most of the changes mediated by *Trichoderma* were related to general stress and regulation of biological processes. The Kurmi cultivar might have higher tolerance to microbe attacks due to the expression of genes involved in the biosynthesis of flavonol glycosides and a clade of GLP-defensins unique to quinoa. These genes are thus candidates for selection of quinoa cultivars with higher resistance to microbe attacks.

## Additional files

**Additional file 1: Table S1. Quinoa genes significantly upregulated in the cultivar Kurmi but not in Real.** The table shows genes that were significantly upregulated in the Kurmi cultivar when treated with either BOL-12 or T22. The family of GLPs is highlighted in light green. The flavonoid biosynthetic pathway is highlighted in light purple. The numbers indicate averages of CPM values for each treatment (n = 3).

**Additional file 2: Table S2. Quinoa genes significantly downregulated in the cultivar Kurmi but not in Real.** Genes observed to be significantly downregulated in the Kurmi cultivar when treated with either BOL-12 or T22 are included. The numbers indicate average of CPM values for every treatment (n = 3). CPM of reference genes are showed at the bottom for transparency.

**Additional file 3: Table S3. Quinoa genes significantly up- and downregulated in the cultivar Real but not in Kurmi.** Genes shown were significantly and consistently upregulated or downregulated in the Real cultivar when treated with either BOL-12 or T22. Highlighted in green we can observe a family of chitinases. Light purple highlights WRKY genes and orange highlights ethylene-responsive genes.

**Additional file 4: Table S4. Singular enrichment analysis of differentially expressed genes in quinoa roots treated with Trichoderma.** For each quinoa-*Trichoderma* interaction, quinoa genes differentially expressed (DE) were annotated for Gene Ontology with Argot2 and then analyzed for singular enrichment analysis with AgriGO2. Stress-related GO-term are highlighted in grey and cell wall-related terms in orange.

**Additional file 5: Table S5. Gene expression assessed by RNA-seq and qRT-PCR.** RNA from quinoa roots treated with *Trichoderma* or mock treated (12 hpi) were analysed by RNA-seq and qRT- PCR in order to determine the correlation of expression levels. Fold change was determined by comparing samples treated with each *Trichoderma* strain against the mock-treated control.

**Additional file 6: Table S6. Gene expression in quinoa shoot and root at 36 hpi with Trichoderma.** Quinoa shoot and root samples were assessed by qRT-PCR after 36 h treatment with *Trichoderma* added to the roots. Gene expression was normalized to the *CqAct2* reference gene. Fold change was determined by comparing samples treated with *Trichoderma* against mock-treated. Significant differences between treatment and control is highlighted in red. ND, not detected.

**Additional file 7: Table S7. Primer sequences of quinoa genes analysed by qRT-PCR.** Forward and reverse primer sequences for qRT-PCR. Primer pairs were designed using Perlprimer aiming for exon-exon borders. *CqAct2* were used as reference genes for normalization of the mRNA abundances and were further verified by the *CqMon1* housekeeping gene.

**Additional file 8: Table S8. Quinoa GLPs significantly upregulated upon treatment with Trichoderma.** Phytozome and NCBI codes for the germin-like proteins significantly upregulated by *Trichoderma* in the Kurmi cultivar. These GLPs belong to a quinoa-specific clade (Figure 6).

**Additional file 9: Table S9. Differentially expressed quinoa genes upon treatment with Trichoderma.** Log2 fold changes, log2 CPM values, P-values and False Discovery Rates for the genes that were differentially expressed in any of the quinoa-*Trichoderma* interactions

## Acknowledgments

We are grateful to Proinpa Institute (Quipaquipani, Bolivia) for the generous donation of quinoa seeds and the Instituto de Investigaciones Farmaco-bioquímicas - UMSA (La Paz, Bolivia) for providing *Trichoderma harzianum* BOL-12QD.

Genome mapping, read count and blastn computations were performed on resources provided by SNIC (Swedish National Infrastructure for Computing) through Uppsala Multidisciplinary Center for Advanced Computational Science (UPPMAX, Uppsala, Sweden) under Project SNIC b2015011/b2016265 along with resources provided by UMSA (Universidad Mayor de Sán Andres) through Centro Nacional de Computación Avanzada para Bioinformática y Genómica (CnCaBo, La Paz, Bolivia).

