## Supplementary File 1 for "Transcriptomic analysis of quinoa reveals a group of germin-like proteins induced by *Trichoderma*"

### Table S1. Quinoa genes significantly upregulated in the cultivar Kurmi but not in Real

The table shows genes that were significantly upregulated in the Kurmi cultivar when treated with either BOL-12 or T22. The family of GLPs is highlighted in light green. The flavonoid biosynthetic pathway is highlighted in light purple. The numbers indicate averages of CPM values for each treatment (n = 3).

| Quinoa gene name | Quinoa gene code | Gene description[a] | Kurmi | | | Real | | |
|---|---|---|---|---|---|---|---|---|
| | | | Ctrl | B12 | T22 | Ctrl | B12 | T22 |
| CqGLP-7 | AUR62025241 | Plant defensin | 0,4 | 13,1 | 4,4 | 14,6 | 40,4 | 30,8 |
| PR-10 | AUR62025514 | Major allergen Api g isoallergen 2 | 0,3 | 11,5 | 1,4 | 3,2 | 7,3 | 8,3 |
| CqGLP-17 | AUR62025235 | Plant defensin | 0,6 | 13,5 | 3,2 | 13,1 | 24,8 | 26,3 |
| CYP81E8 | AUR62038793 | Cytochrome P450 81E8 | 1,5 | 31,6 | 6,8 | 20,2 | 50,0 | 29,3 |
| COMT1 | AUR62005985 | Caffeic acid 3-O-methyltransferase | 0,9 | 17,2 | 3,4 | 13,6 | 23,4 | 21,4 |
| CHS | AUR62013677 | Chalcone synthase | 3,0 | 51,3 | 12,2 | 41,9 | 83,5 | 75,1 |
| CqGLP-10 | AUR62025242 | Plant defensin | 2,9 | 39,2 | 18,2 | 47,6 | 108,5 | 91,6 |
| CqGLP-2 | AUR62025224 | Plant defensin | 1,9 | 22,0 | 10,8 | 23,7 | 49,3 | 47,5 |
| CqGLP-9 | AUR62028952 | Plant defensin | 0,6 | 6,6 | 4,6 | 13,7 | 28,2 | 22,4 |
| STPS | AUR62038591 | Probable sesquiterpene synthase | 6,8 | 75,3 | 26,4 | 81,3 | 151,1 | 97,6 |
| CqGLP-6 | AUR62025237 | Plant defensin | 2,1 | 22,9 | 12,2 | 36,5 | 89,6 | 61,8 |
| NN | AUR62026572 | Protein of unknown function | 0,5 | 4,9 | 1,3 | 8,4 | 14,1 | 14,6 |
| CqGLP-14 | AUR62025218 | Plant defensin | 1,9 | 19,8 | 15,2 | 32,1 | 71,6 | 73,7 |
| RCE1 | AUR62043544 | NEDD8-conjugating enzyme Ubc12 | 0,7 | 7,1 | 3,1 | 0,4 | 0,9 | 0,7 |
| CqGLP-15 | AUR62025225 | Plant defensin | 3,5 | 34,8 | 26,6 | 41,5 | 101,4 | 91,3 |
| CqGLP-18 | AUR62025236 | Plant defensin | 4,4 | 38,4 | 22,3 | 87,0 | 136,1 | 153,9 |
| CqGLP-21 | AUR62028956 | Plant defensin | 0,6 | 5,0 | 2,4 | 9,0 | 11,5 | 13,1 |
| Chit1 | AUR62033302 | Chitotriosidase-1 | 0,3 | 2,8 | 1,9 | 1,4 | 3,3 | 3,3 |
| CqGLP-12 | AUR62028948 | Plant defensin | 3,6 | 28,8 | 15,2 | 31,5 | 63,5 | 53,5 |
| D4H | AUR62028217 | Deacetoxyvindoline 4-hydroxylase | 11,9 | 74,2 | 39,1 | 87,5 | 152,4 | 123,8 |
| cmbl | AUR62002636 | Carboxymethylenebutenolidase homolog | 1,5 | 8,7 | 7,5 | 32,8 | 33,6 | 41,9 |
| CISZOG1 | AUR62026077 | Cis-zeatin O-glucosyltransferase 1 | 1,9 | 11,2 | 4,3 | 3,7 | 11,0 | 9,8 |
| NN | AUR62028130 | At1g58390 Probable disease resistance | 0,6 | 3,1 | 2,7 | 5,1 | 3,1 | 2,6 |
| CqGLP-3 | AUR62025232 | Plant defensin | 0,4 | 2,3 | 2,4 | 8,2 | 13,1 | 15,9 |
| NN | AUR62040548 | Protein of unknown function | 0,4 | 2,1 | 1,2 | 2,8 | 3,4 | 2,5 |
| CHI3 | AUR62030612 | Probable chalcone-flavonone isomerase 3 | 6,9 | 34,4 | 18,0 | 26,1 | 54,6 | 47,6 |

| | | | | | | | | |
|---|---|---|---|---|---|---|---|---|
| CqGLP-13 | AUR62025219 | Plant defensin | 0,7 | 3,7 | 3,4 | 6,0 | 12,2 | 12,6 |
| CqGLP-1 | AUR62025223 | Plant defensin | 7,6 | 34,7 | 29,4 | 40,0 | 93,6 | 77,0 |
| CqPR4 | AUR62001001 | Pathogenesis-related protein P2 | 6,9 | 30,0 | 15,6 | 38,7 | 68,2 | 54,4 |
| UGT73C5 | AUR62024488 | UDP-glycosyltransferase 73C5 | 1,7 | 7,6 | 3,9 | 5,9 | 13,0 | 9,2 |
| NN | AUR62037762 | Protein of unknown function | 0,5 | 2,1 | 1,7 | 5,1 | 4,0 | 7,0 |
| CqGLP-19 | AUR62028958 | Plant defensin | 10,7 | 41,9 | 43,3 | 67,0 | 144,2 | 118,9 |
| CqGLP-20 | AUR62040806 | Plant defensin | 3,9 | 15,0 | 12,1 | 59,9 | 102,1 | 78,5 |
| BURP5 | AUR62011666 | BURP domain-containing protein 5 | 7,8 | 30,0 | 21,7 | 33,3 | 58,5 | 60,1 |
| NN | AUR62016052 | Protein of unknown function | 0,3 | 1,4 | 1,9 | 1,3 | 0,5 | 0,3 |
| CG5412 | AUR62019002 | UPF0483 protein CG5412 | 2,2 | 8,0 | 5,0 | 7,2 | 10,7 | 9,5 |
| CqGLP-16 | AUR62025234 | Plant defensin | 16,9 | 56,5 | 41,9 | 75,5 | 132,7 | 130,1 |
| NN | AUR62011182 | Protein of unknown function | 0,9 | 2,9 | 2,6 | 2,3 | 1,6 | 2,1 |
| FL | AUR62014672 | Flavonol synthase/flavanone 3-hydroxylase | 2,2 | 6,1 | 6,2 | 7,8 | 11,4 | 12,5 |
| NN | AUR62033227 | At5g57670 | 1,8 | 4,8 | 4,0 | 2,5 | 3,0 | 3,2 |
| PER8 | AUR62007739 | Peroxidase 8 | 1,5 | 3,9 | 3,2 | 6,4 | 4,5 | 4,1 |
| NN | AUR62020367 | Protein of unknown function | 1,5 | 3,9 | 4,0 | 2,9 | 1,8 | 2,5 |
| CHI | AUR62020547 | Chalcone-flavonone isomerase | 27,8 | 68,2 | 43,8 | 63,1 | 100,2 | 100,1 |
| RR9 | AUR62026868 | Response regulator ORR9 | 2,4 | 5,9 | 9,2 | 6,4 | 4,0 | 6,1 |
| NN | AUR62027119 | Protein of unknown function | 5,6 | 13,3 | 9,6 | 29,3 | 38,4 | 52,5 |
| NN | AUR62041108 | Protein of unknown function | 3,6 | 8,5 | 7,4 | 8,8 | 6,7 | 9,4 |
| NN | AUR62001977 | GDSL esterase/lipase At2g23540 | 12,2 | 26,0 | 23,6 | 46,9 | 73,2 | 96,6 |
| NN | AUR62026150 | Protein of unknown function | 4,8 | 10,1 | 8,6 | 14,5 | 14,9 | 20,7 |
| NN | AUR62003568 | Protein of unknown function | 32,4 | 67,1 | 57,7 | 74,2 | 113,8 | 113,6 |
| NN | AUR62006030 | Protein of unknown function | 3,9 | 7,7 | 13,0 | 4,9 | 3,4 | 3,6 |
| CHI3 | AUR62024932 | Probable chalcone-flavonone isomerase 3 | 39,4 | 78,6 | 61,5 | 66,8 | 99,1 | 104,7 |
| CqRSLP 2 | AUR62031352 | Os03g0733400 RICESLEEPER 2 | 11,1 | 21,7 | 19,2 | 22,4 | 23,3 | 35,9 |
| SAUR71 | AUR62001328 | Auxin-responsive protein SAUR71 | 5,1 | 9,9 | 8,3 | 10,0 | 6,4 | 8,3 |
| SPAC644.07 | AUR62018315 | Probable mitochondrial chaperone bcs1 | 12,2 | 23,1 | 18,0 | 13,7 | 24,9 | 22,5 |
| CAR4 | AUR62018204 | Protein C2-DOMAIN ABA-RELATED 4 | 7,7 | 14,4 | 13,6 | 20,8 | 21,6 | 27,9 |
| NN | AUR62023337 | Protein of unknown function | 24,4 | 42,2 | 36,1 | 58,0 | 38,3 | 56,1 |
| GDPD1 | AUR62029437 | GDPD1 | 11,1 | 19,0 | 25,0 | 27,0 | 22,7 | 26,9 |
| CHI | AUR62015655 | Chalcone-flavonone isomerase | 75,8 | 126,7 | 102,3 | 102,7 | 132,9 | 133,2 |
| CDC2 | AUR62033172 | Cell division control protein 2 homolog | 12,6 | 20,8 | 23,4 | 19,3 | 17,7 | 29,3 |

[a] Jarvis et al., 2017

### Table S2. Quinoa genes significantly downregulated in the cultivar Kurmi but not in Real

Genes observed to be significantly downregulated in the Kurmi cultivar when treated with either BOL-12 or T22 are included. The numbers indicate average of CPM values for every treatment (n = 3). CPM of reference genes are showed at the bottom for transparency.

| Quinoa gene name | Quinoa gene code | Gene description[a] | Kurmi | | | Real | | |
|---|---|---|---|---|---|---|---|---|
| | | | Ctrl | B12 | T22 | Ctrl | B12 | T22 |
| *CDC20-1* | AUR62028149 | Cell division cycle 20.1 | 77,3 | 50,1 | 36,1 | 45,8 | 72,7 | 32,3 |
| *CDC20-1* | AUR62005915 | Cell division cycle 20.1 cofactor of APC complex | 52,9 | 34,2 | 24,4 | 32,6 | 45,6 | 23,0 |
| *OLP* | AUR62001811 | Osmotin-like protein | 34,3 | 21,2 | 15,5 | 20,9 | 34,5 | 17,8 |
| *NN* | AUR62036106 | Protein of unknown function | 70,0 | 43,3 | 40,0 | 48,0 | 74,6 | 36,9 |
| *NN* | AUR62009169 | Protein of unknown function | 34,2 | 20,0 | 15,9 | 13,0 | 30,7 | 13,3 |
| *SWEET6B* | AUR62029552 | Bidirectional sugar transporter SWEET6b | 28,7 | 16,5 | 15,2 | 16,7 | 21,5 | 18,7 |
| *PG* | AUR62030002 | Polygalacturonase At1g48100 | 28,3 | 15,4 | 9,7 | 18,8 | 16,9 | 14,7 |
| *WAG1* | AUR62019653 | Serine/threonine-protein kinase WAG1 | 33,0 | 17,3 | 17,6 | 14,9 | 18,3 | 15,2 |
| *WAG1* | AUR62013960 | Serine/threonine-protein kinase WAG1 | 13,1 | 6,8 | 6,0 | 6,9 | 7,8 | 4,3 |
| *LBD41* | AUR62023311 | LOB domain-containing protein 41 | 7,8 | 3,7 | 1,9 | 2,2 | 5,2 | 3,0 |
| *PCKA* | AUR62005878 | Phosphoenolpyruvate carboxykinase [ATP] | 27,9 | 13,1 | 13,6 | 4,2 | 5,0 | 4,3 |
| *LECRKS7* | AUR62028107 | Probable L-type lectin-domain containing receptor kinase S.7 | 5,5 | 2,4 | 3,0 | 2,7 | 3,2 | 3,4 |
| *NN* | AUR62004646 | Protein of unknown function | 8,8 | 3,4 | 4,1 | 3,3 | 0,9 | 2,8 |
| *CqEGlu* | AUR62043172 | Glucan endo-beta-glucosidase | 3,3 | 1,0 | 0,9 | 0,4 | 0,5 | 1,0 |
| *MHB1* | AUR62011380 | Non-symbiotic hemoglobin 1 | 10,4 | 2,9 | 2,9 | 1,5 | 1,9 | 2,4 |
| *PG* | AUR62002459 | Polygalacturonase | 6,1 | 1,3 | 1,7 | 1,5 | 1,9 | 1,1 |
| *CqAct2A* | AUR62014374 | CqActin2 Ref Gene | 806,8 | 787,1 | 539,6 | 710,1 | 724,8 | 687,1 |
| *CqAct2B* | AUR62019116 | CqActin2 Ref Gene | 812,1 | 692,9 | 527,3 | 655,8 | 661,4 | 612,3 |
| *CqAct2C* | AUR62014579 | CqActin2 Ref Gene | 81,8 | 70,0 | 60,9 | 74,7 | 69,6 | 57,0 |
| *CqAct2D* | AUR62039382 | CqActin2 Ref Gene | 141,1 | 131,8 | 112,7 | 130,0 | 113,5 | 91,7 |
| *CqMon1A* | AUR62020295 | CqMon1 Ref Gene | 45,6 | 41,0 | 41,6 | 38,3 | 39,6 | 35,2 |
| *CqMon1B* | AUR62037705 | CqMon1 Ref Gene | 35,2 | 36,3 | 33,2 | 26,8 | 28,3 | 27,0 |

[a] Jarvis et al., 2017

### Table S3. Quinoa genes significantly up- and downregulated in the cultivar Real but not in Kurmi.

Genes shown were significantly and consistently upregulated or downregulated in the Real cultivar when treated with either BOL-12 or T22. Highlighted in green we can observe a family of chitinases. Light purple highlights WRKY genes and orange highlights ethylene-responsive genes.

| Quinoa gene name | Quinoa gene code | Gene description |
|---|---|---|
| OPR3 | AUR62025200 | 12-oxophytodienoate reductase 3 |
| DODA1 | AUR62012347 | 2C5-DOPA dioxygenase extradiol 1 |
| KCS1 | AUR62022664 | 3-ketoacyl-CoA synthase 1 |
| KCS2 | AUR62006329 | 3-ketoacyl-CoA synthase 2 |
| ABCG6 | AUR62025538 | ABC transporter G family member 6 |
| CYP74A | AUR62040103 | Allene oxide synthase chloroplastic |
| nep1 | AUR62038206 | Aspartic proteinase nepenthesin-1 |
| nep2 | AUR62015131 | Aspartic proteinase nepenthesin-2 |
| BAM3 | AUR62018803 | Beta-amylase 3%2C chloroplastic |
| CAMBP25 | AUR62018430 | Calmodulin-binding protein 25 |
| Cht4 | AUR62027403 | Chitinase 4 |
| ATX1 | AUR62000774 | Copper transport protein ATX1 |
| CYP86A8 | AUR62004370 | Cytochrome P450 86A8 |
| CYP86A8 | AUR62022579 | Cytochrome P450 86A8 |
| ATL6 | AUR62000873 | E3 ubiquitin-protein ligase ATL6 |
| HOS3 | AUR62008021 | Elongation of fatty acids protein 3-like |
| EPHX4 | AUR62007573 | Epoxide hydrolase 4 |
| ERF4 | AUR62017052 | Ethylene-responsive transcription factor 4 |
| ERF5 | AUR62018880 | Ethylene-responsive transcription factor 5 |
| ERF9 | AUR62014855 | Ethylene-responsive transcription factor 9 |
| ERF017 | AUR62016128 | Ethylene-responsive transcription factor ERF017 |
| EXO70A1 | AUR62008535 | Exocyst complex component EXO70A1 |
| EXO70B1 | AUR62009665 | Exocyst complex component EXO70B1 |
| XLG1 | AUR62016663 | Extra-large guanine nucleotide-binding protein 1 |
| SKIP30 | AUR62028597 | F-box/kelch-repeat protein SKIP30 |
| CYP75B1 | AUR62008515 | Flavonoid 3'-monooxygenase |

| | | |
|---|---|---|
| CYP75B2 | AUR62021575 | Flavonoid 3'-monooxygenase |
| CYP75A1 | AUR62009892 | Flavonoid 5'-hydroxylase 1 |
| FH8 | AUR62026311 | Formin-like protein 8 |
| At5g24080 | AUR62021884 | G-type lectin S-receptor-like serine/threonine-protein kinase At5g24080 |
| GATA5 | AUR62010402 | GATA transcription factor 5 |
| At5g13200 | AUR62005695 | GEM-like protein 5 |
| GA2OX2 | AUR62011753 | Gibberellin 2-beta-dioxygenase 2 |
| At4g29360 | AUR62020357 | Glucan endo-3-beta-glucosidase 12 |
| GPAT4 | AUR62004368 | Glycerol-3-phosphate 2-O-acyltransferase 4 |
| GPAT4 | AUR62022577 | Glycerol-3-phosphate 2-O-acyltransferase 4 |
| GPAT5 | AUR62012122 | Glycerol-3-phosphate acyltransferase 5 |
| GPAT5 | AUR62040714 | Glycerol-3-phosphate acyltransferase 5 |
| HSF24 | AUR62001930 | Heat shock factor protein HSF24 |
| HHP1 | AUR62013296 | Heptahelical transmembrane protein 1 |
| LAC12 | AUR62012314 | Laccase-12 |
| LAC12 | AUR62022954 | Laccase-12 |
| LHT1 | AUR62023955 | Lysine histidine transporter 1 |
| LHT1 | AUR62031750 | Lysine histidine transporter 1 |
| 1MMP | AUR62007090 | Metalloendoproteinase 1-MMP |
| MLP43 | AUR62043142 | MLP-like protein 43 |
| MYB4 | AUR62015573 | Myb-related protein Myb4 |
| mhkB | AUR62011395 | Myosin heavy chain kinase B |
| CHLN | AUR62000396 | Nicotianamine synthase |
| PAE2 | AUR62002521 | Pectin acetylesterase 2 |
| PER24 | AUR62034604 | Peroxidase 24 |
| PER24 | AUR62036667 | Peroxidase 24 |
| At4g16820 | AUR62018869 | Phospholipase A1-Ibeta2%2C chloroplastic |
| At1g06800 | AUR62008926 | Phospholipase A1-Igamma1 chloroplastic |
| PGIP2 | AUR62024807 | Polygalacturonase inhibitor 2 |
| MJ0612 | AUR62013277 | Probable arogenate/prephenate dehydrogenase |
| CML36 | AUR62026154 | Probable calcium-binding protein CML36 |
| CAF1-11 | AUR62020248 | Probable CCR4-associated factor 1 homolog 11 |
| CAD6 | AUR62018276 | Probable cinnamyl alcohol dehydrogenase 6 |

| | | |
|---|---|---|
| At1g62630 | AUR62025629 | Probable disease resistance protein At1g62630 |
| GATL1 | AUR62016655 | Probable galacturonosyltransferase-like 1 |
| At1g74360 | AUR62021617 | Probable LRR receptor-like serine/threonine-protein kinase |
| RKF3 | AUR62006561 | Probable LRR receptor-like serine/threonine-protein kinase RKF3 |
| At2g30020 | AUR62020159 | Probable protein phosphatase 2C 25 |
| WRKY11 | AUR62020288 | Probable WRKY transcription factor 11 |
| WRKY23 | AUR62039260 | Probable WRKY transcription factor 23 |
| WRKY33 | AUR62026343 | Probable WRKY transcription factor 33 |
| WRKY40 | AUR62030836 | Probable WRKY transcription factor 40 |
| WRKY41 | AUR62010821 | Probable WRKY transcription factor 41 |
| WRKY41 | AUR62019820 | Probable WRKY transcription factor 41 |
| WRKY42 | AUR62021917 | Probable WRKY transcription factor 42 |
| WRKY57 | AUR62003119 | Probable WRKY transcription factor 57 |
| WRKY70 | AUR62029778 | Probable WRKY transcription factor 70 |
| XTH23 | AUR62027741 | Probable xyloglucan endotransglucosylase/hydrolase protein XTH23 |
| EXO | AUR62027794 | Protein EXORDIUM |
| EXL2 | AUR62027796 | Protein EXORDIUM-like 2 |
| LYK5 | AUR62043269 | Protein LYK5 |
| SARD1 | AUR62033803 | Protein SAR DEFICIENT 1 |
| YLS9 | AUR62037049 | Protein YLS9 |
| YLS9 | AUR62044049 | Protein YLS9 |
| RGA4 | AUR62033818 | Putative disease resistance protein RGA4 |
| RPPL1 | AUR62007274 | Putative disease resistance RPP13-like protein 1 |
| PGSIP7 | AUR62042795 | Putative glucuronosyltransferase PGSIP7 |
| PERK11 | AUR62008808 | Putative proline-rich receptor-like protein kinase PERK11 |
| ATL2 | AUR62015272 | RING-H2 finger protein ATL2 |
| ATL2 | AUR62018069 | RING-H2 finger protein ATL2 |
| SCL13 | AUR62021833 | Scarecrow-like protein 13 |
| SCL5 | AUR62009552 | Scarecrow-like protein 5 |
| At3g07070 | AUR62020037 | Serine/threonine-protein kinase At3g07070 |
| At5g01020 | AUR62030734 | Serine/threonine-protein kinase At5g01020 |
| SDR2a | AUR62009713 | Short-chain dehydrogenase reductase 2a |
| HSP23 | AUR62039962 | Small heat shock protein chloroplastic |

| | | |
|---|---|---|
| SYP132 | AUR62036357 | Syntaxin-132 |
| MYB44 | AUR62010308 | Transcription factor MYB44 |
| RAX3 | AUR62012493 | Transcription factor RAX3 |
| TYDC2 | AUR62025133 | Tyrosine/DOPA decarboxylase 2 |
| PUB26 | AUR62010239 | U-box domain-containing protein 26 |
| PUB4 | AUR62003956 | U-box domain-containing protein 4 |
| UGT87A2 | AUR62001891 | UDP-glycosyltransferase 87A2 |
| At3g50280 | AUR62018684 | Uncharacterized acetyltransferase At3g50280 |
| At3g28850 | AUR62034387 | Uncharacterized protein At3g28850 |
| SAP7 | AUR62017364 | Zinc finger A20 and AN1 domain-containing stress-associated protein 7 |
| ZAT12 | AUR62001834 | Zinc finger protein ZAT12 |
| ZAT12 | AUR62009622 | Zinc finger protein ZAT12 |
| ZAT12 | AUR62038383 | Zinc finger protein ZAT12 |
| NN | AUR62034001 | 5-epi-aristolochene synthase 3 |
| CqChit1 | AUR62027407 | Acidic endochitinase |
| NN | AUR62010867 | B2 protein |
| NN | AUR62019730 | Basic 7S globulin 2 |
| NN | AUR62011155 | Cucumber peeling cupredoxin |
| CqChit1 | AUR62019040 | Endochitinase |
| NN | AUR62039951 | Hydrophobic seed protein |
| NN | AUR62002452 | Protein of unknown function |
| NN | AUR62003653 | Protein of unknown function |
| NN | AUR62007952 | Protein of unknown function |
| NN | AUR62008152 | Protein of unknown function |
| NN | AUR62008676 | Protein of unknown function |
| NN | AUR62009687 | Protein of unknown function |
| NN | AUR62010274 | Protein of unknown function |
| NN | AUR62011712 | Protein of unknown function |
| NN | AUR62012437 | Protein of unknown function |
| NN | AUR62012922 | Protein of unknown function |
| NN | AUR62012928 | Protein of unknown function |
| NN | AUR62016627 | Protein of unknown function |
| NN | AUR62017310 | Protein of unknown function |

| | | |
|---|---|---|
| NN | AUR62018946 | Protein of unknown function |
| NN | AUR62020224 | Protein of unknown function |
| NN | AUR62021162 | Protein of unknown function |
| NN | AUR62022193 | Protein of unknown function |
| NN | AUR62023931 | Protein of unknown function |
| NN | AUR62024933 | Protein of unknown function |
| NN | AUR62025086 | Protein of unknown function |
| NN | AUR62025343 | Protein of unknown function |
| NN | AUR62027173 | Protein of unknown function |
| NN | AUR62027736 | Protein of unknown function |
| NN | AUR62031561 | Protein of unknown function |
| NN | AUR62032323 | Protein of unknown function |
| NN | AUR62033664 | Protein of unknown function |
| NN | AUR62034721 | Protein of unknown function |
| NN | AUR62038186 | Protein of unknown function |
| NN | AUR62040531 | Thaumatin-like protein 1 |

### Table S4. Singular enrichment analysis of differentially expressed genes in quinoa roots treated with *Trichoderma*

For each quinoa-*Trichoderma* interaction, quinoa genes differentially expressed (DE) were annotated for Gene Ontology with Argot2 and then analyzed for singular enrichment analysis with AgriGO2. Stress-related GO-term are highlighted in grey and cell wall-related terms in orange.

| Interaction | GO term | Description | DE genes | Total genes | p-value | FDR |
|---|---|---|---|---|---|---|
| Kurmi and T22 | - | None | - | - | - | - |
| Kurmi and BOL-12 | GO:0006952 | defense response | 6 | 108 | 5,8E-05 | 0,01 |
| | GO:0009607 | response to biotic stimulus | 5 | 83 | 1,6E-04 | 0,02 |
| Real and BOL-12 | GO:0006073 | cellular glucan metabolic process | 6 | 117 | 2,9E-05 | 0,00 |
| | GO:0044042 | glucan metabolic process | 6 | 117 | 2,9E-05 | 0,00 |
| | GO:0044264 | cellular polysaccharide metabolic process | 6 | 117 | 2,9E-05 | 0,00 |
| | GO:0005976 | polysaccharide metabolic process | 6 | 140 | 7,8E-05 | 0,01 |
| | GO:0044262 | cellular carbohydrate metabolic process | 6 | 174 | 2,6E-04 | 0,02 |
| | GO:0043043 | peptide biosynthetic process | 10 | 599 | 1,0E-03 | 0,03 |
| | GO:0034645 | cellular macromolecule biosynthetic process | 22 | 1977 | 7,3E-04 | 0,03 |
| | GO:0044271 | cellular nitrogen compound biosynthetic process | 21 | 1907 | 1,0E-03 | 0,03 |
| | GO:0071554 | cell wall organization or biogenesis | 5 | 144 | 7,9E-04 | 0,03 |
| | GO:0043604 | amide biosynthetic process | 10 | 599 | 1,0E-03 | 0,03 |
| | GO:0009059 | macromolecule biosynthetic process | 22 | 1979 | 7,4E-04 | 0,03 |
| | GO:0006412 | translation | 10 | 591 | 9,3E-04 | 0,03 |
| | GO:0006518 | peptide metabolic process | 10 | 613 | 1,2E-03 | 0,04 |
| | GO:0043603 | cellular amide metabolic process | 10 | 622 | 1,4E-03 | 0,04 |
| Real and T-22 | GO:0080090 | regulation of primary metabolic process | 80 | 983 | 4,9E-12 | 0,00 |
| | GO:0060255 | regulation of macromolecule metabolic process | 81 | 1001 | 4,9E-12 | 0,00 |
| | GO:2001141 | regulation of RNA biosynthetic process | 78 | 933 | 2,4E-12 | 0,00 |

| | | | | | |
|---|---|---|---|---|---|
| GO:0009889 | regulation of biosynthetic process | 79 | 961 | 4,0E-12 | 0,00 |
| GO:0006355 | regulation of transcription, DNA-templated | 78 | 933 | 2,4E-12 | 0,00 |
| GO:0010556 | regulation of macromolecule biosynthetic process | 79 | 961 | 4,0E-12 | 0,00 |
| GO:1903506 | regulation of nucleic acid-templated transcription | 78 | 933 | 2,4E-12 | 0,00 |
| GO:0051252 | regulation of RNA metabolic process | 78 | 935 | 2,6E-12 | 0,00 |
| GO:0031326 | regulation of cellular biosynthetic process | 79 | 961 | 4,0E-12 | 0,00 |
| GO:0031323 | regulation of cellular metabolic process | 80 | 983 | 4,9E-12 | 0,00 |
| GO:2000112 | regulation of cellular macromolecule biosynthetic process | 79 | 961 | 4,0E-12 | 0,00 |
| GO:0010468 | regulation of gene expression | 80 | 974 | 3,1E-12 | 0,00 |
| GO:0019219 | regulation of nucleobase-containing compound metabolic process | 78 | 947 | 4,9E-12 | 0,00 |
| GO:0019222 | regulation of metabolic process | 81 | 1003 | 5,4E-12 | 0,00 |
| GO:0051171 | regulation of nitrogen compound metabolic process | 79 | 968 | 5,7E-12 | 0,00 |
| GO:0016567 | protein ubiquitination | 19 | 84 | 5,0E-11 | 0,00 |
| GO:0032446 | protein modification by small protein conjugation | 19 | 86 | 7,7E-11 | 0,00 |
| GO:0036211 | protein modification process | 133 | 2135 | 3,0E-10 | 0,00 |
| GO:0006464 | cellular protein modification process | 133 | 2135 | 3,0E-10 | 0,00 |
| GO:0044260 | cellular macromolecule metabolic process | 251 | 4736 | 5,5E-10 | 0,00 |
| GO:0043412 | macromolecule modification | 134 | 2195 | 8,6E-10 | 0,00 |
| GO:0097659 | nucleic acid-templated transcription | 78 | 1126 | 1,1E-08 | 0,00 |
| GO:0006351 | transcription, DNA-templated | 78 | 1126 | 1,1E-08 | 0,00 |
| GO:0032774 | RNA biosynthetic process | 78 | 1130 | 1,2E-08 | 0,00 |
| GO:0019438 | aromatic compound biosynthetic process | 86 | 1293 | 1,4E-08 | 0,00 |
| GO:0043170 | macromolecule metabolic process | 266 | 5308 | 2,0E-08 | 0,00 |
| GO:1901362 | organic cyclic compound biosynthetic process | 88 | 1347 | 2,1E-08 | 0,00 |
| GO:0034654 | nucleobase-containing compound biosynthetic process | 80 | 1208 | 4,6E-08 | 0,00 |

| | | | | | |
|---|---|---|---|---|---|
| GO:0044237 | cellular metabolic process | 288 | 5911 | 6,7E-08 | 0,00 |
| GO:0018130 | heterocycle biosynthetic process | 84 | 1308 | 8,3E-08 | 0,00 |
| GO:0006468 | protein phosphorylation | 102 | 1739 | 3,2E-07 | 0,00 |
| GO:0070647 | protein modification by small protein conjugation or removal | 19 | 141 | 3,7E-07 | 0,00 |
| GO:0016310 | phosphorylation | 102 | 1845 | 3,8E-06 | 0,00 |
| GO:0050789 | regulation of biological process | 88 | 1533 | 3,8E-06 | 0,00 |
| GO:0050794 | regulation of cellular process | 87 | 1511 | 3,8E-06 | 0,00 |
| GO:0044267 | cellular protein metabolic process | 146 | 2859 | 4,1E-06 | 0,00 |
| GO:0008152 | metabolic process | 438 | 10087 | 1,0E-05 | 0,00 |
| GO:0065007 | biological regulation | 89 | 1610 | 1,3E-05 | 0,00 |
| GO:0005976 | polysaccharide metabolic process | 16 | 140 | 2,4E-05 | 0,00 |
| GO:0071704 | organic substance metabolic process | 310 | 7010 | 2,8E-05 | 0,00 |
| GO:0006796 | phosphate-containing compound metabolic process | 109 | 2110 | 3,0E-05 | 0,00 |
| GO:0006793 | phosphorus metabolic process | 109 | 2117 | 3,4E-05 | 0,00 |
| GO:0006073 | cellular glucan metabolic process | 14 | 117 | 4,5E-05 | 0,00 |
| GO:0044264 | cellular polysaccharide metabolic process | 14 | 117 | 4,5E-05 | 0,00 |
| GO:0044042 | glucan metabolic process | 14 | 117 | 4,5E-05 | 0,00 |
| GO:0009987 | cellular process | 310 | 7066 | 4,6E-05 | 0,00 |
| GO:0046348 | amino sugar catabolic process | 6 | 22 | 6,9E-05 | 0,00 |
| GO:0016998 | cell wall macromolecule catabolic process | 6 | 22 | 6,9E-05 | 0,00 |
| GO:1901071 | glucosamine-containing compound metabolic process | 6 | 22 | 6,9E-05 | 0,00 |
| GO:0044036 | cell wall macromolecule metabolic process | 6 | 22 | 6,9E-05 | 0,00 |
| GO:1901072 | glucosamine-containing compound catabolic process | 6 | 22 | 6,9E-05 | 0,00 |
| GO:0006026 | aminoglycan catabolic process | 6 | 22 | 6,9E-05 | 0,00 |
| GO:0006030 | chitin metabolic process | 6 | 22 | 6,9E-05 | 0,00 |

| | | | | | |
|---|---|---|---|---|---|
| GO:0006032 | chitin catabolic process | 6 | 22 | 6,9E-05 | 0,00 |
| GO:0006040 | amino sugar metabolic process | 6 | 22 | 6,9E-05 | 0,00 |
| GO:0044271 | cellular nitrogen compound biosynthetic process | 98 | 1907 | 8,1E-05 | 0,00 |
| GO:0016070 | RNA metabolic process | 83 | 1575 | 1,1E-04 | 0,00 |
| GO:0044238 | primary metabolic process | 292 | 6719 | 1,1E-04 | 0,00 |
| GO:0019538 | protein metabolic process | 158 | 3387 | 1,4E-04 | 0,00 |
| GO:1901136 | carbohydrate derivative catabolic process | 7 | 35 | 1,5E-04 | 0,00 |
| GO:0006022 | aminoglycan metabolic process | 6 | 30 | 4,4E-04 | 0,01 |
| GO:0044249 | cellular biosynthetic process | 117 | 2500 | 6,5E-04 | 0,01 |
| GO:1901576 | organic substance biosynthetic process | 117 | 2516 | 8,0E-04 | 0,01 |
| GO:0034645 | cellular macromolecule biosynthetic process | 95 | 1977 | 8,2E-04 | 0,01 |
| GO:0009059 | macromolecule biosynthetic process | 95 | 1979 | 8,5E-04 | 0,01 |
| GO:0032501 | multicellular organismal process | 11 | 105 | 8,9E-04 | 0,01 |
| GO:0044262 | cellular carbohydrate metabolic process | 15 | 174 | 9,3E-04 | 0,01 |
| GO:0006950 | response to stress | 38 | 650 | 1,1E-03 | 0,01 |
| GO:0010467 | gene expression | 96 | 2025 | 1,2E-03 | 0,02 |
| GO:0006633 | fatty acid biosynthetic process | 9 | 82 | 1,8E-03 | 0,02 |
| GO:0016053 | organic acid biosynthetic process | 14 | 173 | 2,4E-03 | 0,03 |
| GO:0072330 | monocarboxylic acid biosynthetic process | 9 | 86 | 2,5E-03 | 0,03 |
| GO:0044283 | small molecule biosynthetic process | 16 | 216 | 3,1E-03 | 0,04 |
| GO:0055114 | oxidation-reduction process | 93 | 2040 | 3,8E-03 | 0,04 |
| GO:0044703 | multi-organism reproductive process | 8 | 75 | 3,8E-03 | 0,04 |
| GO:0044706 | multi-multicellular organism process | 8 | 75 | 3,8E-03 | 0,04 |
| GO:0009875 | pollen-pistil interaction | 8 | 75 | 3,8E-03 | 0,04 |
| GO:0008037 | cell recognition | 8 | 75 | 3,8E-03 | 0,04 |

| | | | | | |
|---|---|---|---|---|---|
| GO:0048544 | recognition of pollen | 8 | 75 | 3,8E-03 | 0,04 |
| GO:0009856 | pollination | 8 | 75 | 3,8E-03 | 0,04 |
| GO:0006979 | response to oxidative stress | 18 | 266 | 4,6E-03 | 0,05 |

### Table S5. Gene expression assessed by RNA-seq and qRT-PCR.

RNA from quinoa roots treated with *Trichoderma* or mock treated (12 hpi) were analysed by RNA-seq and qRT-PCR in order to determine the correlation of expression levels. Fold change was determined by comparing samples treated with each *Trichoderma* strain against the mock-treated control.

| Method | Gene | Code | Kurmi | | Real | |
|---|---|---|---|---|---|---|
| | | | BOL-12 | T22 | BOL-12 | T22 |
| | | | FC(log2) | FC(log2) | FC(log2) | FC(log2) |
| qRT-PCR* | *CqCat2* | *AUR62040809* | 0,78 | 0,71 | 1,09 | 1,35 |
| | *CqEDS* | *AUR62001194* | 0,88 | 0,79 | 0,82 | 0,62 |
| | *CqGLP1* | *AUR62025223* | 4,14 | 4,10 | 1,71 | 1,26 |
| | *CqGLP10* | *AUR62025242* | 14,60 | 8,84 | 2,00 | 1,79 |
| | *CqHSP90* | *AUR62031424* | 1,39 | 2,08 | 1,28 | 2,44 |
| | *CqMon1* | *AUR62020295* | 1,01 | 1,01 | 0,97 | 1,02 |
| | *CqMyc2* | *AUR62018713* | 1,02 | 1,26 | 1,07 | 1,86 |
| | *CqPER39* | *AUR62034603* | 1,14 | 1,56 | 0,54 | 0,61 |
| | *CqPR1C* | *AUR62027040* | 0,91 | 1,24 | 0,32 | 0,50 |
| | *CqRAP2.3* | *AUR62018057* | 0,76 | 0,73 | 0,74 | 0,90 |
| | *CqW33* | *AUR62006298* | 2,33 | 3,52 | 2,36 | 4,35 |
| RNA-seq | *CqCat2* | *AUR62040809* | 0,78 | 0,60 | 1,31 | 1,51 |
| | *CqEDS* | *AUR62001194* | 0,93 | 1,07 | 0,98 | 0,94 |
| | *CqGLP1* | *AUR62025223* | 4,59 | 3,88 | 2,34 | 1,93 |
| | *CqGLP10* | *AUR62025242* | 13,51 | 6,26 | 2,28 | 1,92 |
| | *CqHSP90* | *AUR62031424* | 2,08 | 3,15 | 2,27 | 4,51 |
| | *CqMon1* | *AUR62020295* | 0,90 | 0,91 | 1,04 | 0,92 |
| | *CqMyc2* | *AUR62018713* | 0,89 | 0,90 | 1,94 | 2,99 |
| | *CqPER39* | *AUR62034603* | 0,84 | 1,94 | 0,59 | 0,47 |
| | *CqPR1C* | *AUR62027040* | 1,72 | 1,68 | 0,17 | 0,33 |
| | *CqRAP2.3* | *AUR62018057* | 0,82 | 0,44 | 1,43 | 1,36 |
| | *CqW33* | *AUR62006298* | 1,79 | 4,11 | 3,55 | 7,61 |
| Correlation treatment dependent ($R^2$) | | | 0,90 | 0,86 | 0,85 | 0,92 |
| Correlation ($R^2$) | | | 0,848 | | | |

| | |
|---|---|
| Pearson correlation coefficient | 0,921 |

* Normalized to *CqAct2*

### Table S6. Gene expression in quinoa shoot and root at 36 hpi with *Trichoderma.*

Quinoa shoot and root samples were assessed by qRT-PCR after 36 h treatment with *Trichoderma* added to the roots. Gene expression was normalized to the CqAct2 reference gene. Fold change was determined by comparing samples treated with *Trichoderma* against mock-treated. Significant differences between treatment and control is highlighted in red. ND, not detected.

| Tissue | Gene | Code | Kurmi | | | | Real | | | |
|---|---|---|---|---|---|---|---|---|---|---|
| | | | BOL-12 | | T22 | | BOL-12 | | T22 | |
| | | | FC(log2) | p-value | FC(log2) | p-value | FC(log2) | p-value | FC(log2) | p-value |
| Shoot | *CqCat2* | *AUR62040809* | -0,04 | 0,97 | -0,61 | 0,00 | -0,21 | 0,33 | 0,40 | 0,29 |
| | *CqEDS* | *AUR62001194* | 0,44 | 0,53 | 0,27 | 0,54 | 0,03 | 0,95 | 0,16 | 0,72 |
| | *CqGLP1* | *AUR62025223* | 0,95 | 0,04 | 0,01 | 0,95 | 1,31 | 0,03 | -0,14 | 0,57 |
| | *CqGLP10* | *AUR62025242* | 1,12 | 0,03 | 0,82 | 0,27 | 1,77 | 0,01 | 0,51 | 0,24 |
| | *CqHSP83a* | *AUR62031424* | -0,02 | 0,93 | 1,17 | 0,01 | -0,44 | 0,24 | 1,31 | 0,01 |
| | *CqMon1* | *AUR62020295* | 0,11 | 0,46 | 0,08 | 0,22 | 0,04 | 0,85 | -0,24 | 0,30 |
| | *CqMyc2* | *AUR62018713* | -0,02 | 0,88 | 0,19 | 0,21 | 0,04 | 0,86 | -0,76 | 0,13 |
| | *CqPER39* | *AUR62034603* | ND | - | ND | - | ND | - | ND | - |
| | *CqPR1C* | *AUR62027040* | ND | - | ND | - | ND | - | ND | - |
| | *CqRAP2.3* | *AUR62018057* | 0,07 | 0,62 | -0,55 | 0,30 | -0,04 | 0,76 | 0,21 | 0,25 |
| | *CqW33* | *AUR62006298* | 0,18 | 0,60 | 1,41 | 0,00 | 0,63 | 0,04 | 0,67 | 0,43 |
| Root | *CqCat2* | *AUR62040809* | -0,47 | 0,01 | 0,08 | 0,63 | -0,43 | 0,10 | -0,22 | 0,18 |
| | *CqEDS* | *AUR62001194* | -0,19 | 0,20 | 1,31 | 0,05 | -0,27 | 0,56 | 1,33 | 0,04 |
| | *CqGLP1* | *AUR62025223* | 0,63 | 0,29 | 1,41 | 0,14 | 1,02 | 0,13 | 1,55 | 0,09 |
| | *CqGLP10* | *AUR62025242* | 1,01 | 0,04 | 1,59 | 0,03 | 1,54 | 0,15 | 2,65 | 0,01 |
| | *CqHSP83a* | *AUR62031424* | -0,70 | 0,01 | -0,01 | 0,99 | -0,53 | 0,03 | -0,15 | 0,64 |
| | *CqMon1* | *AUR62020295* | 0,15 | 0,17 | -0,18 | 0,61 | -0,15 | 0,33 | 0,25 | 0,11 |
| | *CqMyc2* | *AUR62018713* | -0,37 | 0,04 | 0,25 | 0,50 | -0,07 | 0,83 | 0,24 | 0,66 |
| | *CqPER39* | *AUR62034603* | 0,30 | 0,54 | 0,22 | 0,67 | -0,63 | 0,20 | -1,37 | 0,02 |
| | *CqPR1C* | *AUR62027040* | 0,39 | 0,34 | 0,56 | 0,55 | 0,48 | 0,10 | -0,55 | 0,39 |
| | *CqRAP2.3* | *AUR62018057* | -0,70 | 0,01 | -0,27 | 0,28 | -0,59 | 0,26 | 0,12 | 0,59 |
| | *CqW33* | *AUR62006298* | 0,09 | 0,81 | 1,41 | 0,00 | -0,20 | 0,61 | 0,60 | 0,03 |

### Table S7. Primer sequences of quinoa genes analysed by qRT-PCR.

Forward and reverse primer sequences for qRT-PCR. Primer pairs were designed using Perlprimer aiming for exon-exon borders. *CqAct2* were used as reference genes for normalization of the mRNA abundances and were further verified by the *CqMon1* housekeeping gene.

| Quinoa gene | *C. quinoa* gene code | Putative encoding gene description | | Primer forward | Primer reverse | Product | PCR eff. |
|---|---|---|---|---|---|---|---|
| *CqAct2* | *AUR62014374* | *actin2* | Reference gene | TACCACAGGTATCGTGCTTGACTC | GATCACGTCCGGCAAGATCC | 113 bp | 1,89 |
| *CqMon1A*[a] | *AUR62020295* | *monensin sensitivity 1* | Reference gene | AAGGATCATCTGACCATAAAGC | TCGTGTCAAGTTAGTTCGGG | 145 bp | 1,98 |
| *CqCat2* | *AUR62040809* | *catalase 2* | Defense-related | CCAGGAGTGAGATATAGATCATGGG | CCCAAAGATTTATCCGCCTGAG | 145 bp | 2,03 |
| *CqEDS* | *AUR62001194* | *enhanced disease susceptibility 1* | Defense-related | TTTGTCAGCTTGTTCATCGT | GTCCTGCATATCTTTCTTCCC | 125 bp | 1,96 |
| *CqPR1C* | *AUR62027040* | *Pathogenesis-related protein 1C* | Defense-related | TGTTTCATTGTCATAACCCTAGCC | ACTGTATGTTCACAACACCCAC | 117 bp | 1,93 |
| *CqGLP1* | *AUR62025223* | *Germin-like protein 1* | Defensin | GCATTACAACTCACTCCTACC | CTTCATCCGCATAACTTCCT | 123 bp | 1,93 |
| *CqGLP10* | *AUR62025242* | *Germin-like protein 10* | Defensin | ATACCAACAACACCGCACAC | GTAAGTTCCCAGCAAATAAAGCAG | 179 bp | 1,97 |
| *CqHSP83* | *AUR62031424* | *heat shock protein 83* | Stress-related | ATTCGGTGTTGGTTTCTACTC | CCAAGTATTCCAACTGATCTTCC | 199 bp | 1,92 |
| *CqMyc2* | *AUR62018713* | *Myc2 transcription factor* | Defense-related | GGAAGTGAAGGAGACGAGAA | CAACCCAGCATACTCCGAAA | 102 bp | 2,00 |
| *CqWRKY33* | *AUR62006298* | *WRKY DNA-binding protein 33* | Defense-related | TCCTTTACACCTGAGACATCCT | ACTGTTCTGTTACCATACCCTGAC | 126 bp | 1,93 |
| *CqPER39* | *AUR62034603* | *Peroxidase 39* | Defense-related | TTGTGATTTGTAATGCAGGTGG | CTCGAGGGCAACTCTTATGG | 149 bp | 2,03 |
| *CqRAP2.3* | *AUR62018057* | *Ethylene response factor related to AP 2.2* | Stress-related | ATGATGAGTATGGGATTCTACGG | CCCAAGCAAAGATTCAAGGT | 122 bp | 2,01 |

[a] The primer pair matches 100% with the *CqMon1B* (AUR62037705) gene sequence which shares 95% nucleotide sequence identity with *CqMon1A*.

### Table S8. Quinoa GLPs significantly upregulated upon treatment with *Trichoderma*.

Phytozome and NCBI codes for the germin-like proteins significantly upregulated by *Trichoderma* in the Kurmi cultivar. These GLPs belong to a quinoa-specific clade (Figure 6).

| Gene name | Phytozome code | NCBI code | Peptide size (aa) |
|---|---|---|---|
| CqGLP-1* | AUR62025223 | XP_021750106.1 | 208 |
| CqGLP-2 | AUR62025224 | XP_021750088.1 | 208 |
| CqGLP-3 | AUR62025232 | XP_021750260.1 | 208 |
| CqGLP-6 | AUR62025237 | XP_021750078.1 | 208 |
| CqGLP-7 | AUR62025241 | XP_021750095.1 | 208 |
| CqGLP-9 | AUR62028952 | XP_021714979.1 | 439** |
| CqGLP-10* | AUR62025242 | XP_021750069.1 | 193 |
| CqGLP-12 | AUR62028948 | XP_021714976.1 | 208 |
| CqGLP-13 | AUR62025219 | XP_021750074.1 | 208 |
| CqGLP-14 | AUR62025218 | XP_021750075.1 | 208 |
| CqGLP-15 | AUR62025225 | XP_021750080.1 | 208 |
| CqGLP-16 | AUR62025234 | XP_021750050.1 | 208 |
| CqGLP-17 | AUR62025235 | XP_021750126.1 | 208 |
| CqGLP-18 | AUR62025236 | XP_021750060.1 | 208 |
| CqGLP-19 | AUR62028958 | XP_021714981.1 | 208 |
| CqGLP-20 | AUR62040806 | XP_021754444.1 | 208 |
| CqGLP-21 | AUR62028956 | XP_021714980.1 | 208 |
| Average size (aa) of quinoa GLPs | | | 207 |

* Genes selected for qRT-PCR analysis

** Discarded from the average measurement due to the presence of putative introns, which might reveal an annotation error
